# Transcriptomic and proteomic analysis of quiescent epimastigotes as a resource for investigating *Trypanosoma cruzi* persistence

**DOI:** 10.1101/2025.09.09.675056

**Authors:** Francisco Olmo, Amy Hesketh, Hage Hayat, Jean-Marc Monneuse, Martin C. Taylor, Fernanda C. Costa, Célie DaSilva, Frédéric Bequet, Fanny Escudié, Derry Mercer, Adrien Saliou, Eric Chatelain, John M. Kelly, Josephine Abi-Ghanem

## Abstract

Chagas disease is caused by infection with the protozoan parasite *Trypanosoma cruzi.* Despite triggering a strong immune response, infections are typically life-long and can result in severe cardiac and/or digestive tract pathology. Current drugs have limited efficacy, and treatment failure is a common outcome. Eliminating a non-replicating *T. cruzi* sub-population that can persist after therapy has been a key challenge for the drug-development community. Here, we describe the transcriptome and proteome profiles of quiescent epimastigote forms of the parasite isolated from exponentially growing cultures on the basis of reduced turnover of transiently-induced red fluorescent protein. This quiescent sub-population was characterised by down-regulation of genes/proteins involved in translation, metabolism, mitochondrial function and DNA replication, and by up-regulation of proteins that promote exit from the cell-cycle in other organisms. These data represent a resource that can be exploited to dissect the mechanistic basis of quiescence and to refine the drug-development screening cascade.

---

The insect-transmitted protozoan *Trypanosoma cruzi* causes Chagas disease, the most serious parasitic infection in Latin America, affecting 6-7 million people^1,2^. In humans, the initial infection is characterised by a patent parasitaemia and widespread dissemination in organs and tissues. Acute stage infections normally manifest as a short-term (2-8 weeks), mild, febrile condition, although myocarditis or meningoencephalitis can develop in some cases^3^. The infection is then controlled by a vigorous adaptive immune response, although parasites are not eliminated, and the infection is life-long^1^. Approximately 30% of individuals subsequently develop chronic heart disease^4–6^ and/or debilitating digestive tract pathology^7^, symptoms that can take years to become apparent. The current drugs, benznidazole and nifurtimox, have limited efficacy, and toxicity can prevent patients from completing the long-dosing regimens (typically 60-90 days)^8,9^. There is an international effort to develop new Chagas disease drugs led by consortia involving academic, commercial and not-for-profit partners. This has led to the identification of several pre-clinical candidates^10–12^, although these have yet to progress to the patient. There are few prospects of a useable vaccine within the foreseeable future.

*T. cruzi* is an obligate intracellular parasite capable of invading any nucleated mammalian cell. Infection is mediated by metacyclic trypomastigotes, the flagellated non-replicative life-cycle stage that is transmitted by triatomine bugs. Within host cells, these differentiate into the smaller ovoid amastigote forms, which contain a much-reduced flagellum and divide by binary fission, reaching numbers >100 in some cell types^13^. Amastigote then further differentiate into bloodstream trypomastigotes, which are also flagellated and non-replicative. These subsequently egress from host cells, and propagate the infection by invading other cells, or through uptake in the bloodmeal of the triatomine vector. It is clear, however, that the *T. cruzi* life-cycle has added complexity not recognised in the classical version. Following invasion, trypomastigotes undergo an initial asymmetric cellular division prior to amastigogenesis, resulting in discard of their flagellum and rapid entry of flagellar proteins into the MHC class I processing pathway^14^. Intracellular replication of amastigotes is asynchronous, with a range of different morphologies co-existing within infected cells, including some that may be intermediate forms^15^. Finally, there is mounting evidence that amastigotes can exit from the cell-cycle and enter a non-replicative state.

Reports of potential quiescent stages in the *T. cruzi* life-cycle include stress-induced cell-cycle arrest^16^, spontaneous dormancy^17^, the detection of drug-tolerant non-replicative amastigotes^18^, and evidence for reduced proliferation during chronic stage infections^13^. The role(s) of these non-replicative/quiescent forms during the parasite life-cycle remain unknown, and it is unclear whether single or distinct mechanisms underpin the phenotype(s). Furthermore, the extent to which quiescence is spontaneous or induced has not been resolved. Of particular concern in terms of treatment, is the possibility that non-proliferative parasites may have enhanced drug-tolerance, analogous to “dormant” bacteria, which often display reduced antibiotic susceptibility^19^. In the case of benznidazole, the small number of amastigotes that persist in mice after treatment are in a non-replicative state^18^. *In vitro,* such persisters are viable and retain an ability to re-enter the cell-cycle and proliferate^20^. Understanding the molecular basis of quiescence and the resulting biochemical adaptations has important implications for Chagas disease therapeutics. For example, it could aid the design of drug classes with an enhanced ability to kill non-replicative parasites and inform the optimisation of treatment regimens using currently available drugs.

Two major factors have constrained our ability to characterise homogeneous quiescent *T. cruzi.* First, there are no known molecular markers that identify parasites with this specific phenotype. Second, most evidence suggests that non-replicative amastigotes represent only a small percentage of the total intracellular population. Attempts to exploit retention of cell tracer dyes to identify and then isolate quiescent parasites have been confounded by the inhibitory effects that such dyes can have on parasite proliferation^13^. To circumvent these issues, we took an alternative strategy by focussing on epimastigotes, the flagellated insect-form of the parasite. We engineered epimastigotes so that tightly-regulated inducible-expression of a red fluorescent protein (RFP) was under the control of tetracycline, or its more lipophilic analogue doxycycline. When these inducers are removed, RFP expression is blocked and replicating parasites return to a fluorescence-negative state. Under these conditions, we hypothesised that quiescent parasites should remain fluorescent relative to those that are proliferative. Here, we describe the generation and purification of such parasites, and their proteomic and transcriptomic profiling.

## Results

### Generation of engineered *T. cruzi* lines to explore quiescence

The key to fractionating a *T. cruzi* population that encompasses different growth phenotypes is robust markers that distinguish between replicating and slow/non-replicating parasites. Here, we exploited *T. cruzi* doxycycline/tetracycline-inducible expression. We reasoned that parasites induced to express RFP, which then entered a quiescent state, would retain the red fluorescent phenotype when the inducer was removed. The approach was based on using the TcINDEX-RFP parasite line^21^ (a derivative of the CL Brener clone), which contains an *RFP* gene integrated into a non-transcribed ribosomal RNA spacer, downstream of T7 promoter and a tetracycline operator (*TetO)* sequence (Fig. 1a). We modified our original system to ensure RFP expression was tightly regulated, by transfecting parasites with the pLEW2X plasmid which contains two copies of the T7 RNA polymerase and Tet repressor (*TetR*) genes, modified to incorporate nuclear localisation signals, as indicated (Fig. 1a) (Methods). Addition of tetracycline or doxycycline to the culture promotes release of TetR from the *TetO*, enabling high-level transcription of *RFP* (Fig. 1b, Extended Data Fig. 1a,b). When the inducer is subsequently removed, RFP expression is blocked, and parasites lose red fluorescence through protein turnover and the dilution effect of replication. To establish the degree to which T7 promoter-driven transcription had been silenced in the absence of tetracycline/doxycycline, we used flow cytometry to assess an epimastigote population that had not been exposed to the inducer. Crucially, no RFP-positive parasites were detected (Fig. 1b, Extended Data Fig. 1 c,d).

**Fig. 1.**
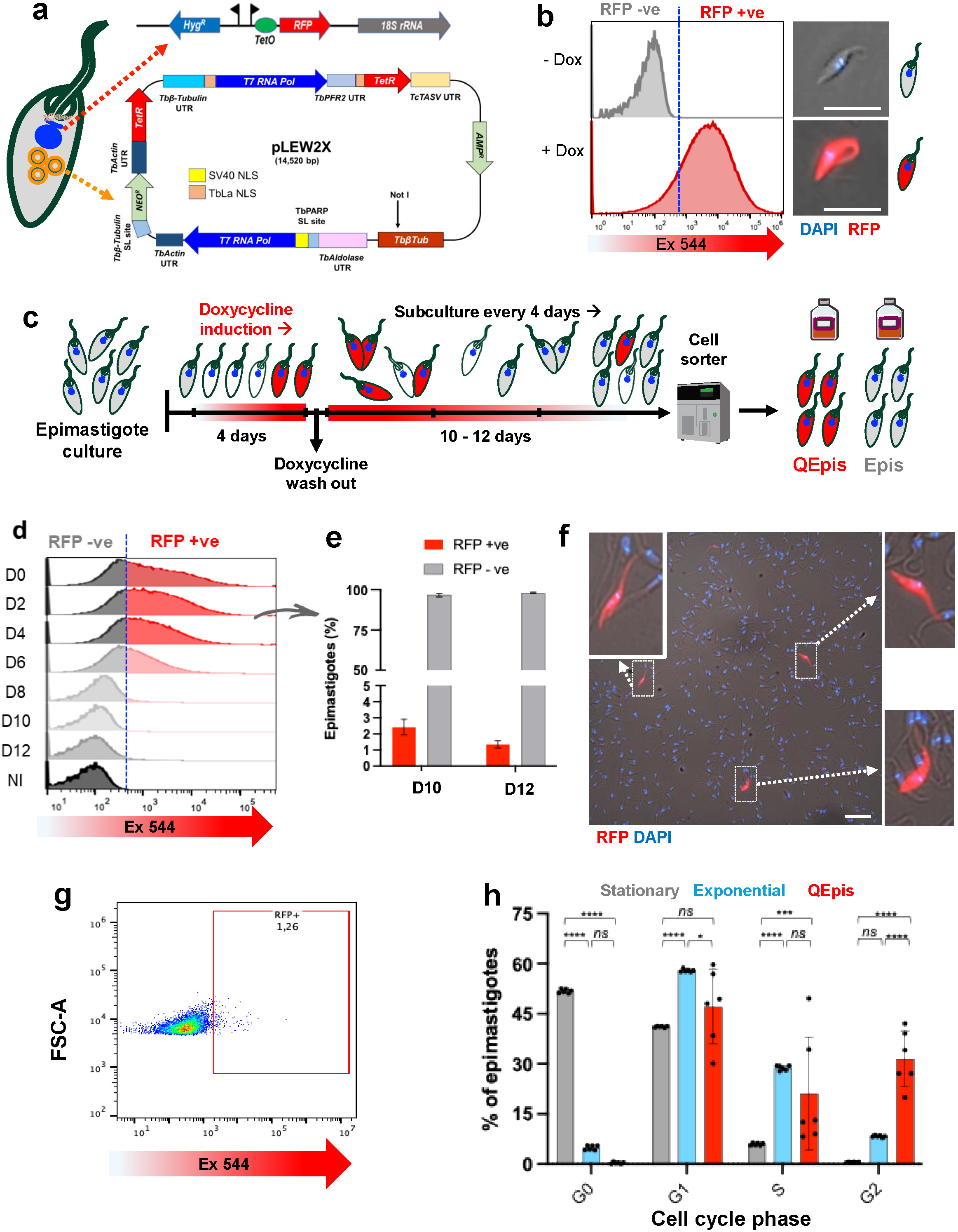
Isolation of quiescent *T. cruzi* epimastigotes (QEpis). **a**, Genetic background of the experimental parasite line. Upper schematic shows the chromosomal locus of the *RFP* integration site in a non-transcribed ribosomal spacer region of the TcINDEX-RFP cell line^21^ downstream of a T7 promoter (flag) and a *TetO* sequence. The cell line was transfected with construct pLEW2X, which contains two copies of the T7 RNA polymerase (*T7 RNA pol*) and tetracycline repressor (*TetR*) genes, modified by insertion of nuclear localisation signals (NLS) as indicated, and selected by growth in G418 (Methods). **b**, Flow cytometry plots of doxycycline induced and non-induced TcINDEX-RFP-pLEW2X epimastigote populations. The RFP+ve/-ve threshold is indicated by a blue dotted line. Adjacent images (right) show representative epimastigotes captured by epifluorescence microscopy. Scale bars = 10 µm. Ex544 (RFP, red), Ex355 (DNA, blue, DAPI staining). **c,** Schematic showing the strategy used to isolate QEpis. (see Methods for further details). **d**, Flow cytometry plots of an induced epimastigote population at 2-day intervals following doxycycline removal. **e**, Bar chart showing the relative percentages of RFP-ve/+ve epimastigotes 10 and 12 days after removal of doxycycline. The results were generated from 10 replicates of 45-50,000 epimastigotes assessed using an Attune NxT cytometer. **f**, Image (Nikon Eclipse iT2 fluorescence microscope) of an induced epimastigotes population 12 days after the removal of doxycycline. Scale bar = 20 µm. Blue (DNA, DAPI staining); Red (RFP). Expanded images (indicated by white dashed arrows) show parasites that have retained RFP and display normal epimastigote morphology. **g,** Flow cytometry panel showing the separation of RFP-retaining parasites (red box, 1.26%) from an RFP-induced culture, after 12 days growth in the absence of doxycycline (as in **c**). **h,** Bar chart illustrating the cell cycle phase distribution of different epimastigote populations. Grey, stationary phase epimastigotes after 3 weeks in culture (Extended Data Fig. X); blue, exponentially growing epimastigotes; red,QEpis). Results are shown as mean ±SD. Two-way ANOVA followed with Turkey’s multiple comparison test ****p < 0.0001.

TcINDEX-RFP-pLEW2X epimastigote cultures were induced to express RFP (Fig. 1c), the doxycycline was then removed, and parasites maintained under exponential growth conditions through regular passage. The population exhibited a steady reduction in red fluorescent parasites, so that 10-12 days after doxycycline removal, the proportion of the parasite population that retained red fluorescence had declined to 1-2% (Fig. 1d-g). We inferred that these parasites, which were motile and displayed a typical epimastigote morphology (Fig. 1f), had entered a non- or low-replicative state, despite optimal growth conditions, and provisionally defined them as quiescent epimastigotes (QEpis). When *T. cruzi* epimastigotes grow in culture, they enter a “stationary phase” where replication stalls, once parasite numbers reach a threshold that leads to nutrient depletion. In this state, parasites are also refractory to doxycycline-induced RFP expression (Extended Data Fig. 1e-g). To compare QEpis with this non-replicative epimastigote form, we assessed their cell cycle profiles. Stationary phase parasites exhibited a predominant G0/G1 cell cycle arrest, with few epimastigotes detected in S or G2 phase, a profile that was rapidly reversible under conditions that promote exponential growth (Fig. 1h, Extended Data Fig. 2). Minimal numbers of QEpis were detected in G0, with the population divided between G1 (50%), S (20%) and G2 (30%) phases. Therefore, QEpis and stationary phase epimastigotes display distinct cell cycle profiles. The QEpi profile was more heterogeneous than that of either stationary or exponentially growing epimastigotes (Fig. 1h), suggesting that QEpis may encompass a more diverse population.

To further investigate the nature of the QEpi population, we separated RFP-retaining and non-retaining epimastigotes by fluorescence-activated cell sorting (FACS) after exposure to an 18-hour pulse of EdU (5-ethynyl-2’-deoxyuridine), a thymidine analogue that is incorporated into replicating DNA (Fig. 2a-d). 40% of non-fluorescent parasites were EdU+ve, and by inference, had been in S-phase during at least some point of the exposure period (Fig. 2b,d). In contrast, only ∼10% of the QEpi population incorporated EdU (Fig. 2c,d). To determine if this reflected a general slow growth profile across the RFP-retaining population, or resulted from some of the parasites re-entering the cell-cycle, we monitored cumulative growth immediately after cell sorting. Within 3 days, non-RFP retaining control cultures were >4 times as dense as the QEpi culture. However, both parasite populations then displayed similar proliferation rates, whilst maintaining the time-lag (Fig. 2e). The most parsimonious explanation is that freshly isolated QEpis are predominantly non-replicative, but the state is transient, and parasites can gradually re-enter the cell-cycle and then undergo normal exponential growth. The transient nature of the quiescent phenotype was demonstrated by examining EdU incorporation by RFP-retaining parasite population 3 days after they had been sorted (Fig. 2e,f).

**Fig. 2.**
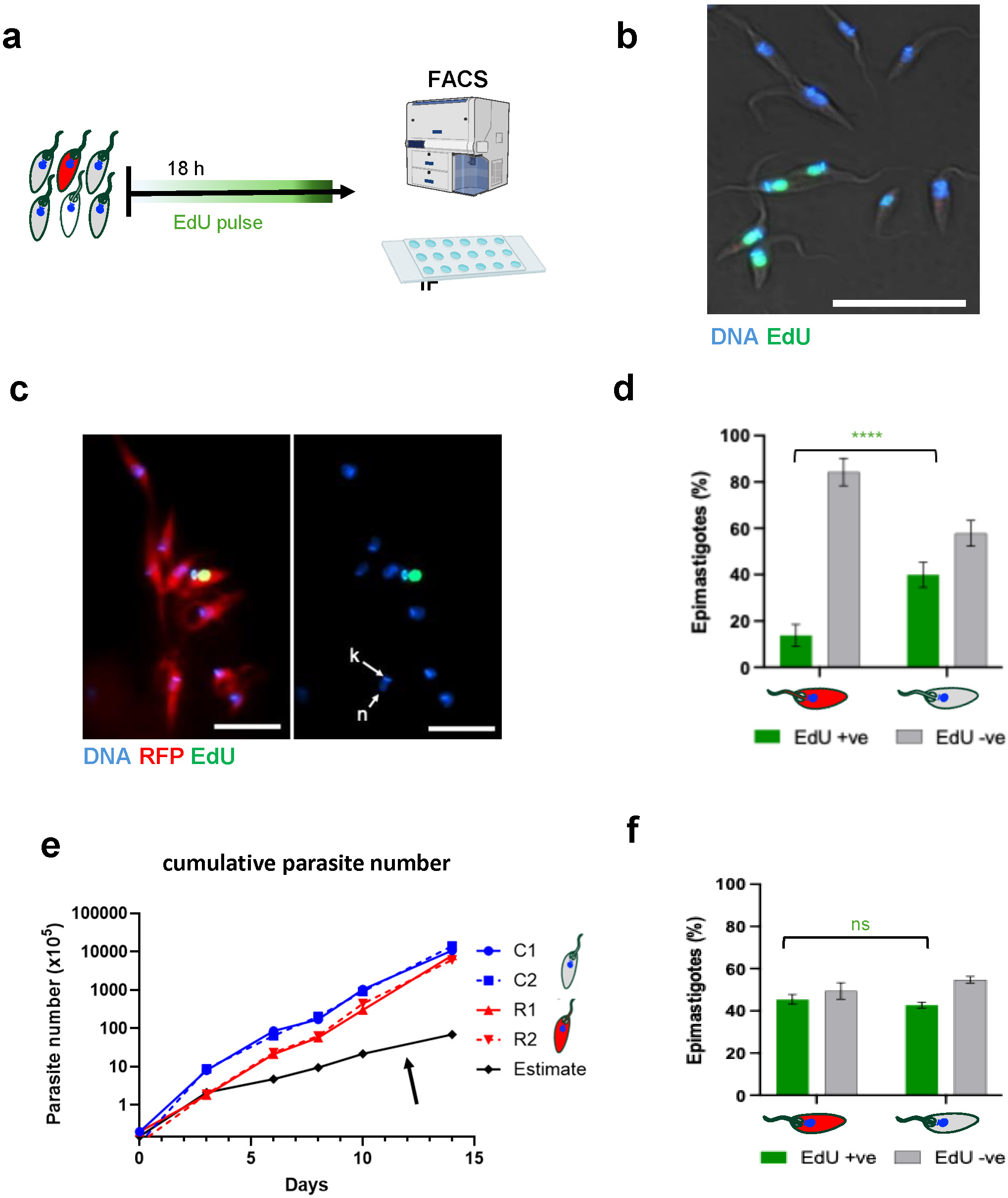
Assessing the replicative status of the QEpi population. **a**, Schematic showing the procedure used to assess replicative status based on EdU incorporation. An RFP-induced culture was maintained in the absence of doxycycline for 11 days (Fig. 1c) and then exposed to EdU for a further 18 hours. After fixing with paraformaldehyde, the QEpis were purified by cell sorter (Methods) and imaged by fluorescence microscopy (IF). **b**, Image of RFP-ve epimastigotes treated as above and purified by FACS. DNA (blue) was stained with DAPI. EdU+ve parasites, which were in S-phase during the exposure period, can be identified by green staining (or turquoise when on a red background). Scale bar = 10 μm. **c**, Images of RFP+ve QEpis (red) after purification by FACS. DNA, blue. An EdU+ve parasite can be identified by green staining (or yellow, when on a red background). White arrows identify nuclear (n) and kinetoplast (k) DNA. Scale bars = 10 μm. **d**, Quantitative analysis of the percentage QEpis (red) and Epis that incorporated EdU during the 18 hour pulse. Results are shown as mean ± SD. One-way ANOVA followed with Šídák’s multiple comparison test **** p <0.0001. **d**, Growth rates of QEpi (R1 and R2) and control epimastigote (C1 and C2) populations. Both sets of parasites were sorted (Methods), sub-cultured at 1 x 10^4^/ml, and their growth monitored for 14 days (counts in triplicate). The black line indicates the predicted growth curve, based on EdU data, if all parasites in the QEpi population had maintained growth at the equivalent reduced rate. **e,** Reversion of QEpi parasites into a replicative state. QEpi parasites and a control group that had lost red fluorescence (generated as in **c**) were sorted, sub-cultured at 1 x 10^4^/ml for 42 hours, and then pulsed with EdU for 18 hours. They were then fixed and sorted (as in **a**) to establish the percentage that had incorporated EdU into replicating DNA; ns, not significant

### Avoiding perturbance of the QEpi transcriptome and proteome profiles

Next, we investigated the nature of the FACS-purified quiescent parasite population by undertaking transcriptomic and proteomic analysis. Given the rarity of QEpi parasites, we sought to avoid lengthy preparation protocols that might perturb RNA/protein expression. We used the commercial reagent ‘CellCover’, which fixes biomolecules within their cellular context without chemical crosslinking. To test the applicability of this procedure, exponentially growing epimastigotes were first used as a control, since preparation of this population requires minimal manipulation. RNA and protein were prepared by two methodologies: one using a standard protocol in which parasites were added directly to the extraction kit, the second in which parasites were “metabolically frozen” using CellCover, and stored at room temperature for 4 hours, and then at 4°C for 96 hours, prior to extraction (Extended Data Fig. 3a) (Methods). A similar number of distinct transcripts were detected with the standard (9,888 genes) and CellCover (9,908 genes) protocols, with <0.1% of transcripts (40 and 60, respectively) uniquely represented (Extended Data Fig. 3b). Only one gene (TcCLB.505941.10, a member of the surface protease GP63 family) was differentially expressed at a significant level between the two populations (log2 fold-change (log2FC) >2.0). At the protein level, a total of 8,010 were detected after preparation with CellCover, and 7,898 after using the standard protocol (Extended Data Fig. 3b). The vast majority (98.5%) of proteins detected were present in both populations. Only 4 proteins were uniquely detected using the standard protocol, in contrast to 116 that were unique to the CellCover set. CellCover may therefore act to protect the proteome. 296 proteins displayed a significant difference in expression levels (log2FC >2) between the two populations.

Taken together, these results suggest that the CellCover procedure does not perturb the transcriptome readout and may even improve protein detection. Since QEpis represent ∼1% of the population at the point of analysis (Fig. 1d-g), and multiple sorting sessions were required to isolate material, we used this protocol in subsequent analysis to avoid perturbation of the transcriptome/proteome profile.

### Transcriptome and proteome profiles of *T. cruzi* life-cycle stages

To provide context for transcriptomic and proteomic data derived from the replicative and non-replicative epimastigote populations (QEpi and Epi, respectively), we compared the ‘omic profiles with those of intracellular amastigotes (a proliferative form) and extracellular trypomastigotes (a non-proliferative form). Using optimized isolation strategies to obtain highly enriched, clearly defined populations (Methods), we detected a total of 9,344 ± 316 distinct transcripts and 4,687 ± 1,822 proteins across all populations (Fig. 3a), a larger number than reported in previous studies^22–27^. The number transcripts detected was similar across different life-cycle stages. In contrast, up to twice as many proteins were identified in the replicative epimastigote population (Epis) compared to non-replicative (QEpis).

**Fig. 3.**
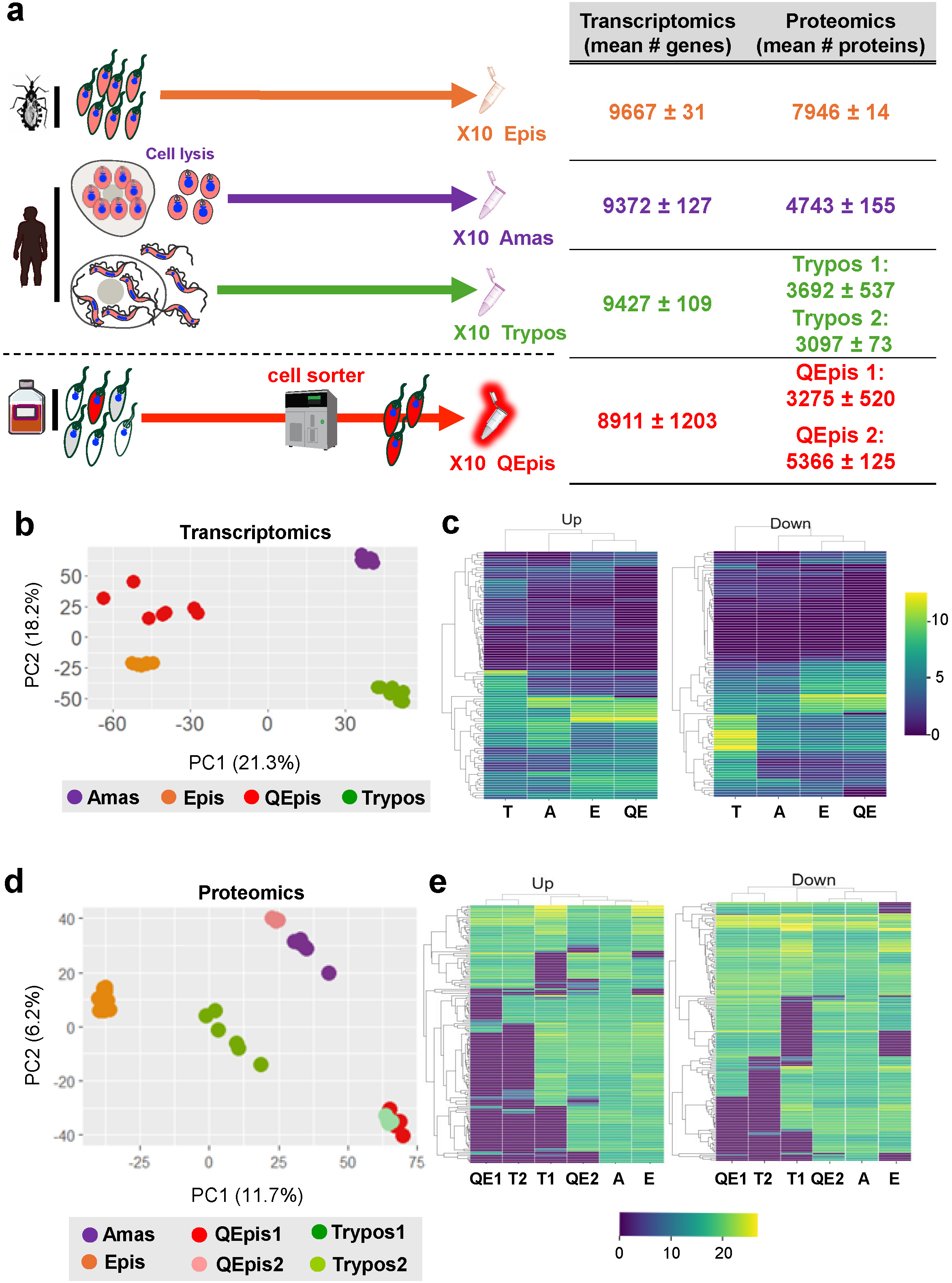
Principal component analysis (PCA) and heatmap visualization of global transcriptomic and untargeted mass spectrometry data. **a**, Number of expressed genes and proteins detected in different *T. cruzi* life-cycle stages. The schematic (left) identifies the four life-cycle stages analysed (see Methods for further details). The data presented represent the mean number of genes and proteins detected in 10 replicates. **b,** PCA plot of transcriptomic data derived from the different parasite life-cycle stages: T (trypomastigotes), A (amastigotes), E (epimastigotes), and QE (quiescent epimastigotes). **c,** Heatmap illustrating the most significant differentially expressed genes categorized as up-regulated or down-regulated across the life-cycle stages. Differential expression analysis was performed using *limma*^46^. Samples were grouped based on their corresponding parasite population and the median of the express values were plotted. Parasite populations are shown in columns and organized by hierarchical clustering. Pixel intensity, ranging from yellow (high) to blue (low) represents the log2FC transformed and normalised expression levels. The percentage values on the axes indicate the proportion of variance explained by each principal component. **d**, PCA plot of proteomic data. Analysis splits the QEpi and Trypo populations into sub-groups 1 and 2 as indicated. **e,** Heatmap showing up-regulated and down-regulated protein expression, organised as outlined above.

Principal component analysis (PCA) of the transcriptome data from amastigotes (Amas), trypomastigotes (Trypos), and epimastigotes (Epis and QEpis), revealed distinct profiles for each population (Fig. 3b,c). However, the QEpis and Epis did share a large number of differentially expressed genes compared with other life-cycle stages. (Fig. 3c). Only 336 genes were up-regulated at an RNA level when the Epi population was compared to QEpis, with 60 down-regulated (log2FC >2; Supplementary Table 1), many of which encoded surface antigens in both cases. Interestingly, transcripts encoding each of the subunits of the mini-chromosome maintenance complex (MCM 2-7), a heterohexamer helicase that is essential for both the initiation and elongation of DNA replication, were down-regulated in QEpis (Extended Data Fig. 4a). In *T. cruzi*, MCM subunits are depleted during metacyclogenesis, as epimastigotes differentiate to metacyclic trypomastigotes, and transition to a state of replicative arrest^28^.

PCA of the proteomes identified a greater degree of clustering into sub-populations, with both trypomastigotes and the QEpis separating into two distinct groups (Trypos1 and 2; QEpis1 and 2) (Fig. 3d,e). The Trypo2 and QEpi1 sub-populations over-lapped, implying related proteomic profiles. Compared to Epis, 2,342 proteins were differentially expressed in QEpis1, and 2,124 in Trypos2 (log2FC >2; Supplementary Table 2), of which 85% were shared. This could reflect their non-proliferative phenotype. In contrast, the QEpi2 sub-population displayed more similarity to amastigotes (a replicative form) (Fig. 3d,e), possibly highlighting a transitional switch by the QEpi2 sub-group from a non-replicative to a replicative state, as suggested by the EdU incorporation data (Fig. 2d). Development of the quiescent phenotype therefore appears to be a transient event, with parasites readily able to switch back to a replicative state (Fig. 2e,f). This sub-division, a reflection of the epimastigote population at the point of protein/RNA preparation, suggests that some parasites had re-entered the cell-cycle, but the level of RFP had yet to be reduced below a threshold that prevents selection during cell sorting. Compared to the Epi population, 470 QEpi2 and 583 Amas proteins were differentially expressed (log2FC >2, Supplementary Table 2), of which 57% were shared. We also detected a similar number of proteins in the QEpis1 and Trypos2 populations, and likewise in the QEpis2 and Amas populations. A mean of 7,946 proteins were detected in Epis (Fig. 3a), the most of any life-cycle stage. Of these, 60% and 79% overlapped with the QEpi1 and QEpi2 sub-populations, respectively. 18 proteins were present in both QEpis 1 and 2, but absent from Epis, and 2 proteins were unique to the QEpi 1 proteome, and 3 unique to QEpi 2 (Supplementary Table 3).

A clear trend across all parasite life-cycle forms was that differential expression was more pronounced at a protein than at an RNA level. This is consistent with previous reports and reflects that transcription in the trypanosomatids is polycistronic and control of gene expression is predominantly a post-transcriptional event, with translational regulation having a key role^29–31^.

### Pathway enrichment analysis

Pathway enrichment analysis provided a better understanding of the nature of the QEpi population in comparison to replicating epimastigotes (Epis). With the transcriptome, up-regulation at a pathway level was only observed in the Epi population (Fig. 4a). The functional groups ‘cellular anatomical structure’ and ‘cellular component’ were prominent (in total, 8 terms enriched) indicating changes in the organelle (GO:0031090, GO:0019866, GO:0031967), cellular (GO:0110165) and mitochondrial (GO:0005739) membranes. For the ‘metabolic process’ functional group (5 terms enriched), the focus was small molecule biosynthesis (nucleobase GO:0055086 and carbohydrate GO:1901135). The fact that there were relatively few differences between the populations at the differential RNA expression level (log2FC >1) could be indicative that the quiescent QEpi population is ‘on-hold’ in preparation for switching to a replicative state. At the protein level, the extent of differential expression (based on log2FC >1) was considerably greater. The Epi population displayed a pathway enrichment profile consistent with a replicative state, with more than 50% of the differentially expressed proteins encompassed by ‘Biological Process’ terms, in comparison to the QEpi 1 and 2 populations (Fig. 4b,c). In the case of the QEpi2 profile, up-regulated protein expression could be encompassed by a single enrichment term (cellular anatomical structure). Pathway enrichment comparison of the QEpi1 and QEpi2 populations (Fig. 4d) revealed a number of pathways that were distinct, although these were far fewer than the differences between the replicating and quiescent epimastigote populations (Fig. 4b,c). In the QEpi1 population, a total of 31 terms were enriched compared to QEpi2, 10 of which were related to flagellum motility, axoneme structure and the dynein complex. An overview of the number of shared and unique proteins within the Epi, QEpi1 and QEpi2 populations is shown in the Venn diagram (Fig. 4e).

**Fig. 4.**
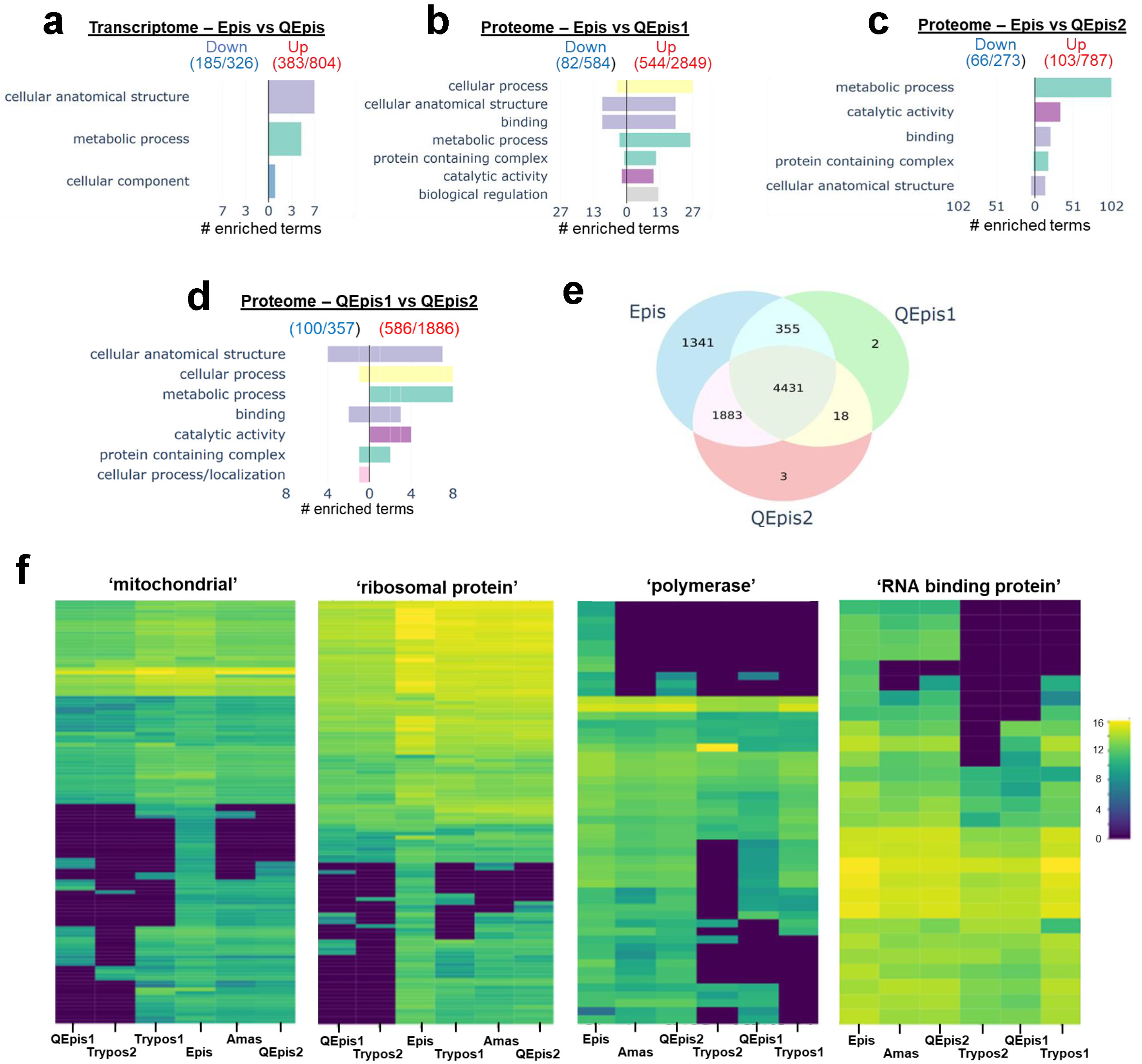
Comparing the Epi and QEpi populations using pathway enrichment analysis. Analysis was conducted using Gene Ontology (GO) terms. A manual grouping was performed to create broader macro terms, allowing for high-level functional assessment. The bar charts (**a-d**) illustrate the pathways enriched in different populations, with the x-axis representing the number of GO terms. As indicated, the right side of the graph identifies up-regulated pathways, and the left side the down-regulated pathways. In brackets, the proteins or genes of unknown function relative to the total number identified (based on log2FC >1) are indicated. For clarity, only macro terms encompassing more than five GO terms are shown. **a,** transcriptomic pathway enrichment comparing Epis and QEpis. **b** and **c,** proteomic pathway enrichment comparing Epis with the QEpi1 and QEpi2 populations, as indicated. **d,** Enrichment analysis of proteins specific to either QEpis1 or QEpis2, as inferred from the Venn diagram. **e**, Venn diagram illustrating shared and unique total protein numbers within the Epi, QEpi1 and QEpi2 populations. **f,** Heatmaps showing expression levels of specific protein groups in different life-cycle populations. The whole proteome data set was interrogated using the search terms ‘mitochondrial’ (118 proteins identified), ‘ribosomal’ (110 proteins identified), ‘polymerase’ (53 proteins identified), and ‘RNA binding protein’ (28 proteins identified). Parasite populations are shown in columns and organized by hierarchical clustering, while individual proteins are displayed in rows, also organized by hierarchical clustering. Pixel intensity represents log expression levels, ranging from yellow (high) to blue (low). A more extensive analysis of these heatmaps is provided in Extended Data Fig. 5-8.

### Classes of protein differentially regulated in quiescent epimastigotes

Compared to the profile of exponentially growing epimastigotes, the QEpi1 data revealed an extensive reduction in expression of proteins associated with a replicative phenotype. In the case of mitochondrial function, for example, proteins involved in kinetoplast DNA replication and maintenance, translation, electron transport, protein assembly/maturation, respiration, energy production, fatty acid metabolism and protein import were widely depleted (Fig. 4f, Extended Data Fig. 5). Six of the down-regulated proteins were members of the mitochondrial carrier family^32^, a class of protein that has a role in transport of metabolites across mitochondrial membranes. Collectively, these data suggest that development of the quiescent QEpi1 phenotype is associated with down-regulation of mitochondrial activity. Also, prominent amongst proteins that exhibited reduced expression in the QEpi1 population in comparison to epimastigotes, were those associated with ribosomes (Fig. 4f). In total, 13 cytosolic and 15 mitochondrial ribosomal proteins were depleted compared with exponentially growing epimastigotes, in addition to several that were linked to the transcription and processing of rRNA (Extended Data Fig. 6). In the QEpi1 parasites, protein synthesis is therefore impacted at several levels by down-regulation of expression.

Quiescence was also linked with lower level expression of enzymes that mediate DNA replication/maintenance and transcription (Fig. 4f). Consistent with the reduced incorporation of EdU (Fig. 2d), these included nuclear DNA polymerases delta, epsilon, sigma and theta, plus several mitochondrial localised DNA polymerases (Extended Data Fig. 7), In addition, subunits of RNA polymerase I, II and III^33^, which mediate transcription of rRNA, protein coding genes, tRNAs and other small RNAs, respectively, were depleted, together with a mitochondrial RNA polymerase and subunits of the MCM complex (Extended Data Fig. 4b). Therefore, the quiescent epimastigotes are characterised by reduced levels of proteins that are central to DNA replication, RNA transcription/processing, and protein synthesis. There was striking overlap in the expression profiles of these proteins between the QEpis1 and Trypos2 populations (Fig. 4f). This may reflect their non-replicative/quiescent phenotypes, and highlights how shared sets of genes may contribute to entry of parasites into this state at different stages of the life-cycle. These data (Fig. 4f) again demonstrate the existence of two distinct trypomastigote groupings (Trypos 1 and 2), as revealed by the PCA analysis (Fig. 3D). As inferred above, the distinct QEpi1 and 2 groupings apparent from the heatmaps, also appear to be representative of two sub-populations, one that is in a quiescent, low/non-replicative state, and one that has, or is, initiating re-entry into the cell-cycle.

In trypanosomatids, RNA binding proteins play a major role in regulating gene expression and cell-cycle control^34^. Analysis of the QEpi1 proteome profile revealed reduced expression of several RNA binding proteins compared with both the Epi and QEpi2 populations (Fig. 4f, Extended Data Fig. 8). These included three distinct mitochondrial RNA binding complex 1 subunit proteins, and two members of the pumilio (Puf) family of RNA binding proteins that are regulators of translation and mRNA stability in many diverse eukaryotes^35^. In *T. cruzi*, one of these proteins (TcPuf3) has a role in controlling mitochondrial morphology and function^36^. Our data also highlighted other RNA binding proteins that might be involved in triggering entry of the parasite into a quiescent state. For example, the transcript encoding an orthologue of the *T. brucei* RNA binding protein TbRBP7 (TcCLB.506565.8), which promotes cell-cycle arrest^37^, was up-regulated in QEpis, relative to replicating epimastigotes (Supplementary Table 1). Analysis of the proteome data also revealed that the RNA binding protein TcRBP6 (TcCLB.506693.30) was up-regulated in QEpis 1 and 2 (Supplementary Table 3). In *T. brucei,* the equivalent protein is a driver of metacyclogenesis, a process that involves differentiation to a non-replicative life-cycle stage^38^. Another protein expressed at increased levels in the QEpi1 and 2 populations, included an orthologue of the YAK kinase family (a CMGC/DYRK protein kinase) (TcCLB.511291.40), which is required for cellular quiescence in yeast^39^ and *Dictyostelium discoideum*^40^, and in *T. brucei*, triggers cell-cycle arrest and entry into a non-replicative state^37^.

## Discussion

Understanding the molecular basis of quiescence in *T. cruzi* has been confounded by technical complexities, and to date, the phenomenon has resisted experimental dissection. Here, we explored if the epimastigote stage of the life-cycle could be exploited as a more tractable model to overcome these issues. With the system we developed (Fig. 1a-c), it was possible to identify a sub-population of epimastigotes (QEpis) that enter into a quiescent state under conditions which normally promote vigorous parasite growth. We also found that QEpis have an ability to revert back to a normal replicative state and continue to proliferate at an exponential rate (Fig. 2e,f). Transient quiescence of *T. cruzi* epimastigotes could have evolved as strategy that prolongs persistence in the insect vector, thereby enhancing transmission. In triatomine bugs, ingested bloodstream trypomastigotes differentiate into epimastigotes which multiply in the mid-gut, migrate to the rectum, and finally differentiate into metacyclic trypomastigotes (Fig. 5). During the next bloodmeal, transmission of these non-replicative infectious parasites to their mammalian host follows their deposition, as part of the excreta, in the vicinity of the bite-wound. The transition of epimastigotes into non-replicative metacyclic trypomastigotes can also occur *in vitro*, for example, when epimastigote cultures reach stationary phase. However, development of this non-replicative state, can lead to differentiation, and is linked to nutrient depletion. Our evidence indicates that this process is distinct from the quiescent phenotype that we report here. First, QEpis were generated under exponential growth conditions (Fig. 1). Second, throughout the process, they displayed an epimastigote-like morphology distinct from that of metacyclic trypomastigotes (Fig.1f). Finally, stationary phase epimastigotes generated in nutrient-depleted medium are predominantly in a state of G0/G1 cell cycle arrest. In contrast, very few QEpi parasite were in G0 (<1% compared to 50% in the stationary phase; Fig. 1h).

**Fig. 5.**
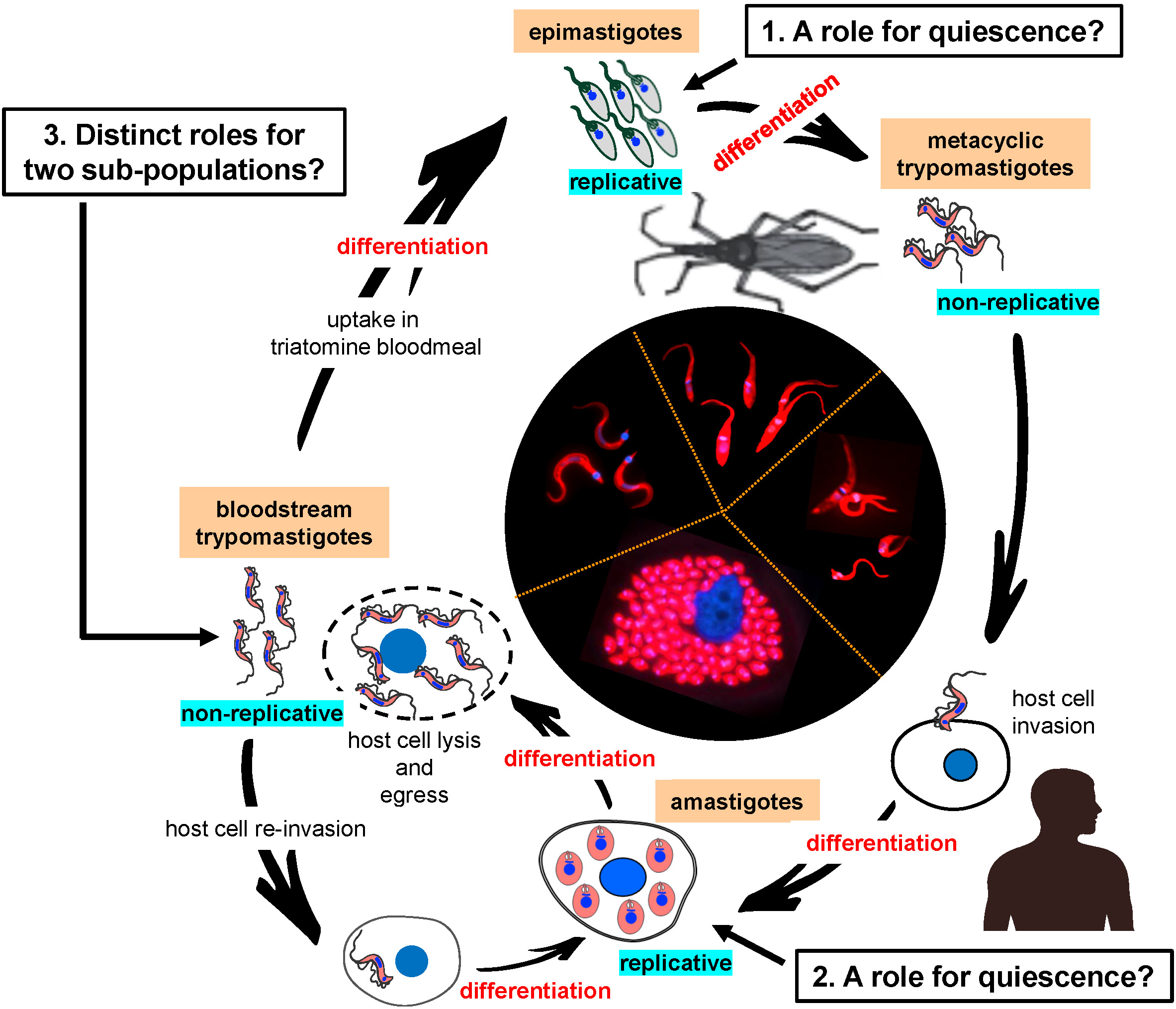
Key questions that should be addressed to more fully understand the *T. cruzi* life-cycle. 1. Is there a role for reversible quiescence in insect stage epimastigotes that may act to prolong infection of the vector and enhance transmission? 2. What is the biological function of quiescence in intracellular amastigotes and does it facilitate long-term persistence and drug tolerance? 3. Are sub-populations of trypomastigotes, as revealed by the proteomic data (Fig. 3d), pre-adapted for distinct roles in the infection/transmission cycle?

It can be inferred from studies utilising incorporation of EdU (Fig. 2) that a sub-population of the QEpi parasites had re-entered a replicative state, although their RFP levels had yet to be depleted below a threshold that avoided selection by cell sorting. Consistent with this, PCA of the proteome data, also revealed two QEpi populations (Fig. 3d). One of these (QEpi2) clustered closely with replicative amastigotes. In contrast, the QEpi1 parasites displayed features more characteristic of a classical quiescent population, with a protein expression profile that reflected widespread down-regulation of key cellular processes such as DNA replication, transcription and protein synthesis (Fig. 4f, Extended Data Fig. 4-8). Several differentially expressed proteins that promote exit from the cell cycle and entry into a non-replicative state in *T. brucei* were also identified. These data therefore provide insight into the nature of a quiescent epimastigote population and identify molecular markers with potential to explore parasite biology and dissect development within the triatomine vector. Importantly, these candidate genes have potential as tools to investigate quiescence in intracellular amastigotes (Fig. 5), either by providing markers for the phenotype, or by identifying potential determinants of the process, features that can be tested experimentally.

Quiescence in *T. cruzi* amastigotes may be a key feature of the parasite life-cycle that facilitates parasite persistence during life-long chronic infections. Analogous to bacterial dormancy, this quiescent phenotype, coupled with an ability to retransition and proliferate under favourable conditions, might also be a significant barrier to drug-mediated elimination^18^. Understanding the molecular drivers of the process could therefore uncover targets that can be exploited for therapeutic purposes, opening doors to novel interventions against Chagas disease. The ability to separate QEpis from proliferative populations (Fig. 1) lays the foundation for future approaches to explore these mechanisms in quiescent amastigotes and to study their functional impact on parasite biology and recrudescence. In the case of bloodstream trypomastigotes, we also identified two distinct sub-populations based on protein expression (Fig. 3d). One possibility is that these groups could correspond to sub-populations that are pre-adapted either to invade other host cells and prolong the infection, or to facilitate transmission after uptake in the bloodmeal of an insect vector (Fig. 5). The differentially expressed genes identified here will therefore provide a means to discriminate between these sub-populations, and to initiate assessment of their biological roles.

In summary, transcriptomic and proteomic analysis of quiescent epimastigotes has provided the first detailed insight into the nature of this poorly characterised feature of the *T. cruzi* life-cycle. The data represent a resource that can be exploited to improve our understanding of parasite biology and will provide a platform to investigate the role of quiescence in long-term persistence within the mammalian host and its possible involvement in drug tolerance.

## Methods

### Parasite and cell culture

*T. cruzi* CL-Brener (DTU-VI) epimastigotes were cultured in RPMI-1640 medium supplemented as previously described^41^. MA104 cells (African green monkey kidney epithelial line) were maintained in complete Minimal Essential Medium (MEM) (Sigma) supplemented with 5% (v/v) heat-inactivated fetal bovine serum (hiFBS, Cytiva), 100 U/ml penicillin, and 100 μg/ml streptomycin. Cells were incubated at 37°C in a 5% CO_2_ atmosphere and sub-cultured (1:5) every 3-4 days.

### *In vitro* infection and trypomastigotes harvesting

For cell monolayer infection, tissue culture trypomastigotes (TCTs) were obtained from previously infected cells. MA104 cell cultures were exposed to TCTs for 18 hours. Extracellular parasites were then removed by washing with PBS, fresh medium added, and cultures incubated for an additional 5-7 days. New extracellular TCTs were isolated by collecting and centrifuging the culture medium at 1600 g. The resulting pellets were resuspended in Dulbecco’s Modified Eagle Medium (DMEM) containing 5% hiFBS and maintained at 37°C for up to 4 hours before use. Motile trypomastigotes were quantified using a hemocytometer.

### Development of the *T. cruzi* inducible system

To enhance silencing of the RFP reporter in the uninduced state, we used a version of the *TetR* gene that included a nuclear localisation signal from the *T. brucei* La protein, derived from plasmid pJ1339^42^ (a gift from Dr Jack Sunter, Oxford Brookes University). The plasmid was digested with EcoRI/AvrII to delete the puromycin acetyltransferase gene and the associated *T. brucei PFR2* 3’-untranslated (3’-UTR) region, then blunt-ended and re-ligated. The resulting plasmid was cut with SpeI, blunt-ended and then digested with NheI to isolate the 4.5 kb band containing the *T7 RNA pol I* and *TetR* genes, both containing an amino-terminal-fused *T. brucei* La protein nuclear localisation signal (Fig. 1a). This fragment was ligated into StuI/NheI cut pLEW13^43^ to create pLEW2X, a construct containing two copies each of the *T7 RNA pol I* and *TetR* genes (Fig. 1a, GenBank Accession # PQ672779). To generate the tetracycline/doxycycline inducible line, *T. cruzi* CL Brener parasites modified to facilitate conditional expression of RFP from a ribosomal locus (the TcINDEX-RFP line^21^) (Figure 1b) were transfected with pLEW2X and selected with 100 μg/mL G418.

### Generation and isolation of quiescent epimastigotes (QEpis)

TcINDEX-RFP epimastigotes transfected with the pLEW2X vector (above) were seeded at 2 × 10^5^ cells/mL in 10 mL of fresh epimastigote medium in non-vented T25 flasks. RFP expression was induced by incubating cultures at 28°C with 1 µg/mL of doxycycline for 96 hours (Fig. 1c). Parasites then underwent two cycles of washing with PBS (4 hours apart) to remove residual doxycycline, and block RFP expression. Following the final wash, parasites were re-seeded at 2 × 10^5^ cells/mL in fresh complete epimastigote medium and maintained in exponential growth for 10 days, with subculturing in fresh medium every 4 days.

To quantify RFP-expressing parasites, 0.5 mL of culture was centrifuged at 3000 g for 5 minutes and fixed with 4% paraformaldehyde (PFA) for 1 hour. Fixed parasites were washed twice with PBS and resuspended in flow cytometry staining buffer (FCSB, Hanks Balanced Salt Solution without calcium/magnesium + 1% BSA). Epimastigotes were then stained with 2 µg/mL Hoechst 33342 for 20 minutes prior to quantification by flow cytometry (Attune NxT Flow Cytometer). Gating was performed using a non-induced culture as control.

For isolation of live RFP-expressing epimastigotes, a similar procedure was used, with the following differences. First, both fixation and Hoechst staining steps were omitted. Second, 8 mL from each flask was centrifuged at 3000 g for 5 minutes and resuspended in 2 mL of CellCover, following a stabilization incubation step at room temperature for 20 minutes. Then, before sorting, 1 mL of FCSB was added. Next, CellCover-protected live parasites were collected by fluorescence-activated cell sorting (BD FACSAria™ II cell sorter) into tubes containing 0.5 mL of CellCover. The sorted population was assessed and quantified by live fluorescence microscopy (Biozero Bz 8000 Fluorescence Microscope) using 10 µL of the collected culture in a disposable cell counting chamber. Finally, parasites were pelleted and washed once with PBS before addition to the lysing solution (PARIS™ kit, Invitrogen) and stored at -80 °C until use.

### Assessing the replication status and viability of Quiescent Epimastigotes (QEpis)

QEpis (RFP-retaining) and control non-RFP-retaining epimastigotes were simultaneously isolated for each replicate by fluorescence-activated cell sorting. Parasites were centrifuged at 3000 g for 5 minutes, resuspended in 1 mL FCSB and sorted directly into Eppendorf tubes containing complete epimastigote medium to minimize perturbation. Each sorted replicate was divided into two groups to assess replicative status and viability.

To determine replicative status, both QEpis and control epimastigotes were incubated for 18 hours with 10 µM EdU (5-ethynyl-2’-deoxyuridine). Epimastigotes were then fixed with 4% PFA at room temperature for 2 hours, washed twice with PBS, and allowed to adhere to 10-well fluorescence slides. EdU incorporation was assessed using a Click-iT Plus EdU AlexaFluor 488 Imaging Kit (Invitrogen) according to the manufacturer’s instructions. Slides were mounted using Vectashield® with DAPI. Images were acquired using an inverted Nikon Eclipse T2i epifluorescence microscope.

In parallel, to determine viability, sorted epimastigotes (QEpis and control) were seeded at 2 × 10^4^ cells/mL and cultured to generate growth curves (Fig. 2e). Statistical analysis was performed using one-way ANOVA followed by Dunnett’s or Tukey’s multiple comparison tests to determine differences in the percentage of EdU-incorporating parasites between populations. Values are expressed as mean ± SD of four replicates. Statistical significance was set at p<0.05.

### Preparation of cell lysates from different life-cycle stages

Epimastigotes were seeded as above, allowed to reach exponential phase by day 4 post-subculture, isolated by centrifugation (1000 g for 5 minutes) and washed twice with PBS. Each replicate was divided into two equal portions: One half was immediately lysed using the PARIS™ kit lysis reagent, flash-frozen on dry ice, and stored at -80°C. The other half was resuspended in CellCover, incubated at room temperature for 4 hours, then stored at 4°C for 96 hours (Extended Data Fig. 1a). These parasites were then pelleted, resuspended in lysis reagent, and stored at -80°C. This allowed the ability of CellCover to preserve RNA and protein content during extended sorting procedures to be assessed.

To generate amastigotes, COLO-N680 cells (a human esophageal squamous cell carcinoma line) were seeded in complete MEM at 80-90% confluency and infected with 2 × 10^6^ TCTs derived as above. After 4 hours, flasks were washed twice with PBS to remove non-internalized parasites and incubated with fresh complete MEM for 72 hours to allow intracellular amastigote replication. Flasks were then washed with PBS, infected cells detached using TrypLE Express (Gibco™), and collected by adding 1 mL HBSS (without calcium/magnesium). They were transferred to a lysing matrix M tube (MP) and homogenized using two 30-second cycles at 5000 m/s in a Precellys 24 homogenizer. The cell lysate containing intact amastigotes was passed through a PD10 desalting column to remove debris. Free amastigotes were concentrated by elution and centrifugation at 3000 g for 5 minutes. Isolated amastigotes were resuspended in PARIS™ kit lysis reagent and stored at -80°C for further analysis.

For trypomastigotes, infected COLO-N680 cell cultures were maintained for 5 days post-infection. Free-swimming trypomastigotes were isolated directly from the culture medium by centrifugation at 3000 g for 5 minutes. The pellet was resuspended in PARIS™ kit lysis reagent as above.

### Transcriptomics

#### RNA extraction

*T. cruzi* samples were shipped on dry ice and thawed on ice upon reception. Samples for transcriptomic analysis were incubated for 10-15 minutes at 37°C, or until the precipitate was dissolved. RNA was then extracted using the PARIS™ kit according to the manufacturer’s recommendations. RNA samples were treated with DNase I and purified using the RNA Clean & Concentrator-5 kit (Zymo Research). RNA samples were then qualified and quantified using the DNF-471 RNA kit on the Fragment Analyzer System (Agilent).

#### Library preparation and sequencing

Sequencing libraries were prepared using the Zymo-Seq RiboFree Total RNA Library kit, following the manufacturer’s recommendation for a total RNA input between 1 and 4 ng. Due to varying RNA extraction yields, inputs for library preparation varied between 0.1 ng and 5 ng. Sequencing libraries were qualified and quantified using the DNF-474 High Sensitivity NGS Fragment kit on the Fragment Analyzer. Libraries were pooled equimolarly and sequenced on the NextSeq2000 System (Illumina) using P3 Reagents in 2x100 cycles (Illumina).

### Proteomics

#### Protein digestion

Paris^TM^ kit lysates were mixed with an equal volume of SP3 lysis buffer (Preomics GmbH) in Covaris AFA tubes. After ultrasonication on a M220 Series ultrasonicator (Covaris) for 90 seconds at 75 W, 20.3% duty factor, 200 bursts per cycle, followed by heating to 95°C. Denatured extract was digested using the SP3-iST kit. Peptide yields were determined using a BCA kit (Pierce™ BCA Protein Assay kits).

#### Mass spectrometry analysis

Peptide analysis was performed on the Ultimate 3500 RS nano LC system coupled to the Exploris 480 mass spectrometer. 350 ng of peptide extracts were injected for each sample. Peptides were separated on a 25 cm EasySpray column (75 µm internal diameter, particle size 2 µm) at a flow rate of 300 nL/minute. After an isocratic step at 2% of solvent B over 3 minutes, the gradient consisted of a first increase of solvent B from 2% to 27.5% over 99 minutes, then from 22% to 40% over 5 minutes. Total run time, including column wash and re-equilibration, was 125 minutes. Tandem mass spectrometry analysis was performed on the Exploris 480 mass spectrometer equipped with an Easy-Spray nanoprobe. The data independent acquisition (DIA) method consisted of a full scan acquired at a resolution of 120,000 from 375 and 1500 m/z (mass to charge ratio) (automatic gain control (AGC) target of 3 x10^6^ ions, or 50 ms maximal injection time) and of 75 DIA windows (isolation 8 m/z, overlap 1 m/z) acquired at 30,000 from 400 to 1008 m/z.

#### Bioinformatic analysis

The quality control of transcriptomic sequencing reads was checked using FASTQC v 0.11.9. Fastp v 0.23.4 was then used to remove low quality reads and to trim any Illumina adapters. Sortmerna v 4.3.6 was used to identify residual rRNA reads remaining after the depletion process, by aligning the reads against the rfam and SILVA rRNA reference database. Filtered reads were then aligned to the corresponding reference genome using STAR v. 2.7.10b. The genome of reference used for this analysis was the *T. cruzi* Esmeraldo like v. 4.2 down-loaded from TriTrypDB. RSeQC package v 5.0.1 was used to assess the mapped reads distribution, coverage uniformity and strand specificity. The aligned reads were used to quantify the number of reads from each genomic feature and to generate the count expression matrix for each gene in each sample, using Salmon v 1.9.0. Count tables from each batch were then merged into a single matrix before the filtering and normalization steps.

In the transcriptomic experiment, four samples had a library size < 0.5 nM, even though they were sequenced. However, these samples showed a lower number of genes. Two of these samples were from the Amas group, with 92 and 222 genes detected, the other two were from the QEpi group, with 3451 and 4496 genes detected. These four samples did not pass the quality control, were considered as outliers, and were removed from the downstream analysis.

#### Extraction and annotation of proteomics

MS spectra was performed using DIA-NN (v. 1.8)^44^ software and Biotracs, an in-house workflow developed with MATLAB (R2021). The DIA-NN spectral library was generated using the proteomes of *Homo sapiens* (id: 9606, down-loaded from Uniprot in February 2023) and *T. cruzi* (id: 5693, down-loaded from Uniprot in April 2023). Trypsin was specified as the enzyme, cleaving after all lysine and arginine residues, and allowing up to two missed cleavages. The N-terminal methionine excision was enabled. Carbamidomethylation of cysteine (+57.021 Da) was specified as a fixed modification and oxidation of methionine (+15.995 Da) and was considered as a variable modification. A False Discovery Rate (FDR) of less than 1% was applied to both peptide spectral matches and protein-based on linear discriminant analysis using a target decoy strategy.

The resulting feature table was used for QC analysis, excluding proteins from the *H. sapiens* database. Robust features present in over 80% of samples per group were retained. Missing features were gap-filled with values from a normal distribution around 1.104. Normalization was performed using the maxLFQ method in DIA-NN^45^.

#### Biostatistical analysis

Differential expression analyses were made for each pair of conditions using the R limma package (v.3.44.3)^46^. Proteins and genes presenting an adjusted p-value <0.05 and an absolute value of log2FC >2, were selected as differentially expressed. Pathway enrichment analyses for transcriptomic data were conducted using the TriTrypDB database [https://tritrypdb.org], and for proteomics using stringDB^48^. For each selected comparison pair, differentially expressed proteins and genes were selected using an absolute value of log2FC >1 and an adjusted p-value <0.05. Gene Ontology and Metabolic Pathway Enrichment were realized for each ontology category (i.e. ’Biological Process’, ’Cellular Component’, ’Molecular Function’), using the ’*T. cruzi* CL Brener Esmeraldo-like’ data, and applying a p-value threshold of 0.05.

#### Data Repository

RNASeq Data are shared via Sequence Read Archive (SRA) N° PRJNA1224747. Proteomics data are shared via ProteomeXchange N° PXD060277

## Acknowledgements

This work was supported by funding from the Drugs for Neglected Diseases initiative (DNDi). DNDi received financial support from: Department for International Development (DFID), UK; Federal Ministry of Education and Research (BMBF) through KfW, Germany; and Médecins sans Frontières (MSF) International.

