## Extended Data Figure legends for "Transcriptomic and proteomic analysis of quiescent epimastigotes as a resource for investigating *Trypanosoma cruzi* persistence"

**Extended Data Fig. 1. Assessing the parameters of doxycycline-induced expression of RFP. a**, Schematic representation of the approach used to induce RFP expression in TcINDEX-RFP-pLEW2X epimastigotes. **b**, Bar chart showing the extent of RFP induction by doxycycline over time using FACS analysis (Methods). Results are the mean of 3 independent replicates. **c**, Fluorescence imaging of exponentially growing epimastigotes cultured in the absence of doxycycline (left) and after 96 hours exposure at 1 μg/mL (right). **d**, Flow cytometry analysis of non-induced epimastigotes (as in **c**). **e**, Schematic representation of the approach used to investigate inducible RFP expression in stationary phase (>3 weeks) epimastigotes. **f**, Flow cytometry plots produced by stationary phase epimastigotes 48 and 96 hours after addition of doxycycline (Ex544) demonstrate that epimastigotes in this state are refractory to inducible RFP expression (upper images). Histograms of cell cycle analysis, performed using Hoechst incorporation (Ex355) (Methods), demonstrates that these parasites remain in G0/G1 arrest throughout the induction period (lower images). **g**, Bar charts of the cell cycle distribution of the induced stationary phase population monitored over 4 consecutive days. Results are the mean of 3 independent replicates.

**Extended Data Fig. 2. G0/G1 arrest of stationary phase cultures is rapidly reversible when parasites are transferred to fresh medium.** When parasites were entering cell cycle arrest (after 1 week in culture), they were either maintained in depleted medium or transferred to fresh medium. With fresh medium, there was a rapid return to an exponentially growing cell cycle profile. Histograms of cell cycle analysis were performed using Hoechst incorporation (Ex355).

**Extended Data Fig. 3. Validation that use of CellCover reagent maintains the integrity of the transcriptome and proteome profiles. a**, Outline of protocols used for generating RNA and protein. Using the standard procedure, epimastigotes (Epis) were pelleted and lysates produced using the PARIS™ kit (Methods). Alternatively, epimastigotes were "metabolically frozen" using CellCover (EpisCC), and stored at 4°C for 96 hours, prior to extraction. Purified RNA and protein were then prepared and used for transcriptome and proteome analysis (Methods). **b**, Comparison of the Epi and EpiCC proteome and transcriptome profiles. Venn diagrams (not to scale) illustrate the total number of transcripts and proteins where identification was methodology-specific. The boxplots show the median number of transcripts and proteins identified in samples from each methodology.

**Extended Data Fig. 4. Expression of the mini-chromosome maintenance (MCM) complex family in different life-cycle populations. a**, Relative expression levels of RNAs that encode subunits 2-7 of the MCM heterohexamer helicase in replicating (Epis) and quiescent (QEpis) epimastigotes. **b**, Heatmap showing protein expression levels of members of the MCM family in different life-cycle populations, as indicated. Pixel intensity represents log expression levels, ranging from yellow (high) to blue (low). MCM subunits 8 and 9 are members of the MCM family, which form a complex, and are involved in DNA repair and replication. Expression of subunits 3 (Q4DGN5_TRYCC), 4(Q4DV49_TRYCC), and 7 (Q4CNA7_TRYCC), which form part of the MCM heterohexamer helicase, was not detected in QEpi1s.

**Extended Data Fig. 5. Heatmap showing expression levels of 'mitochondrial' proteins in different life-cycle populations.** The proteome data sets were interrogated using the search term 'mitochondrial' (118 proteins identified). Parasite populations are shown in columns and organized by hierarchical clustering, while individual proteins are displayed in rows, also organized by hierarchical clustering, with accession numbers shown. Pixel intensity represents log expression levels, ranging from yellow (high) to blue (low). Inset boxes identify proteins where expression is reduced in both QEpi1 and 2 populations compared to epimastigotes. The putative functions of each protein is provided. Note the overlapping profiles of the QEpi1 and Trypo2 populations.

**Extended Data Fig. 6. Heatmap showing expression levels of 'ribosomal proteins' in different life-cycle populations.** The proteome data sets were interrogated using the search term 'ribosomal protein' (110 proteins identified). The heatmap is arranged as described in the legend to Extended Data Fig. 5. Inset boxes highlight proteins where expression is reduced in both QEpi1 and 2 populations compared to epimastigotes, and instances where such reduced expression is restricted to QEpi1s. The putative function of each protein is provided. Note the overlapping profiles of the QEpi1 and Trypo2 populations.

**Extended Data Fig. 7. Heatmap showing expression levels of 'polymerase' proteins in different life-cycle populations.** The proteome data sets were interrogated with the term 'polymerase' (53 proteins identified). The heatmap is arranged as described in the legend to Extended Data Fig. 5. The inset boxes highlight a wide range of DNA and RNA polymerases that are expressed at reduced levels in QEpi1s relative to Epis. The putative function of each of these proteins is provided.

**Extended Data Fig. 8. Heatmap showing expression levels of 'RNA binding proteins' in different life-cycle populations.** The proteome data sets were interrogated with the term 'RNA binding protein' (28 proteins identified). The heatmap is arranged as described in the legend to Extended Data Fig. 5. The inset box identifies 8 putative RNA binding proteins that are expressed at reduced levels relative to epimastigotes.
