## Extended Data Figures 1-8 for "Transcriptomic and proteomic analysis of quiescent epimastigotes as a resource for investigating *Trypanosoma cruzi* persistence"

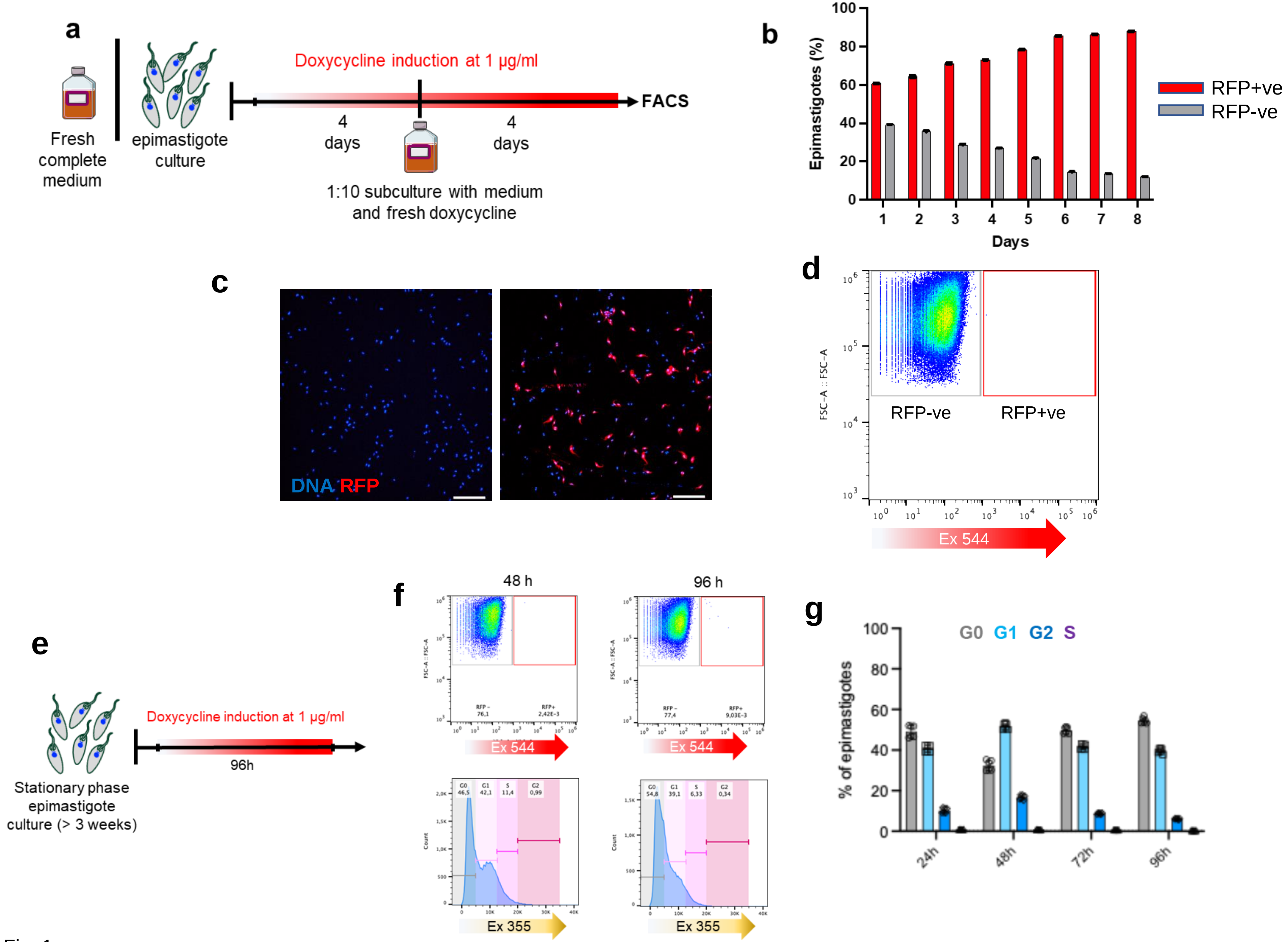


Extended Data Fig. 1

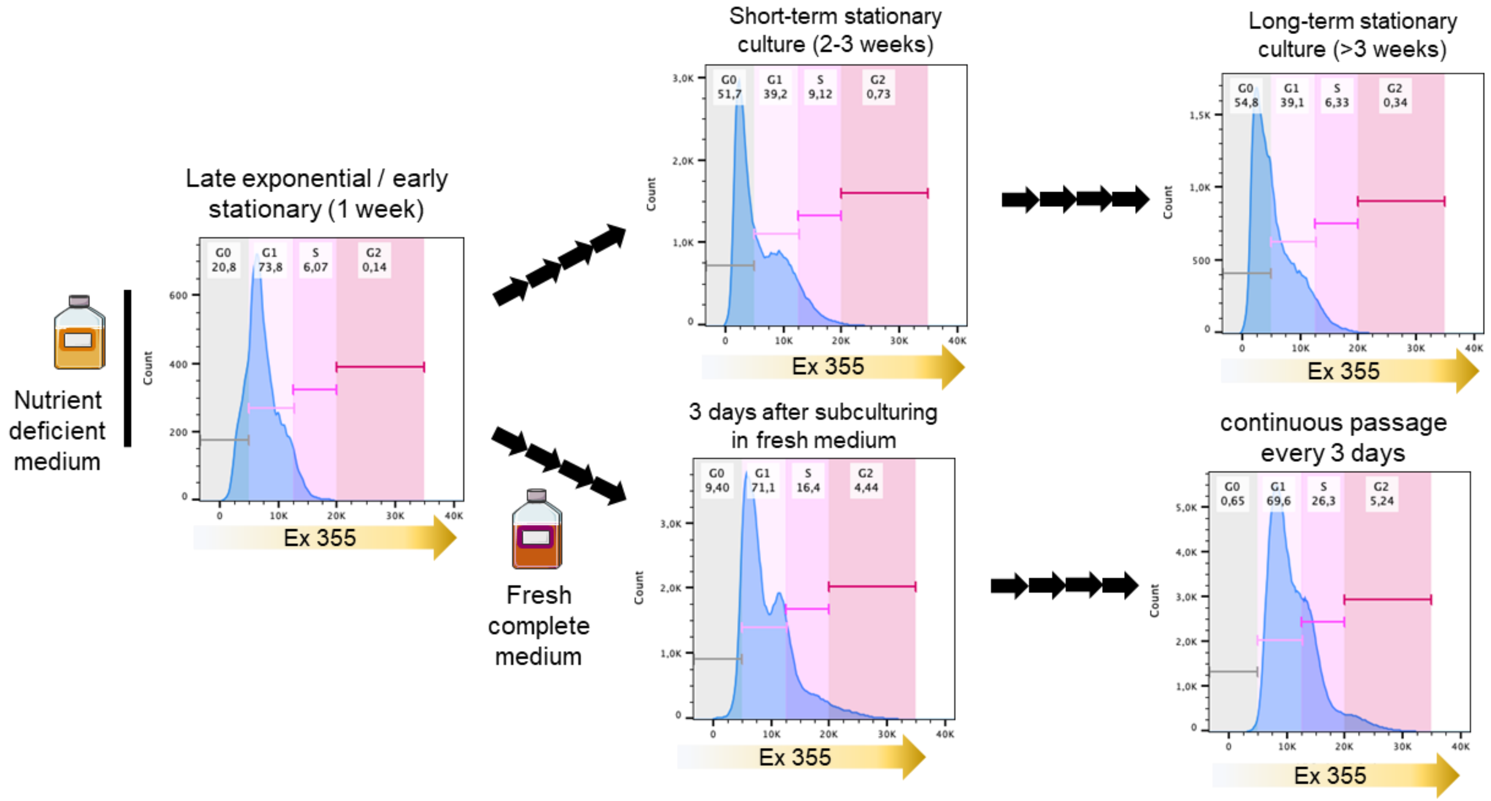


Extended Data Fig. 2

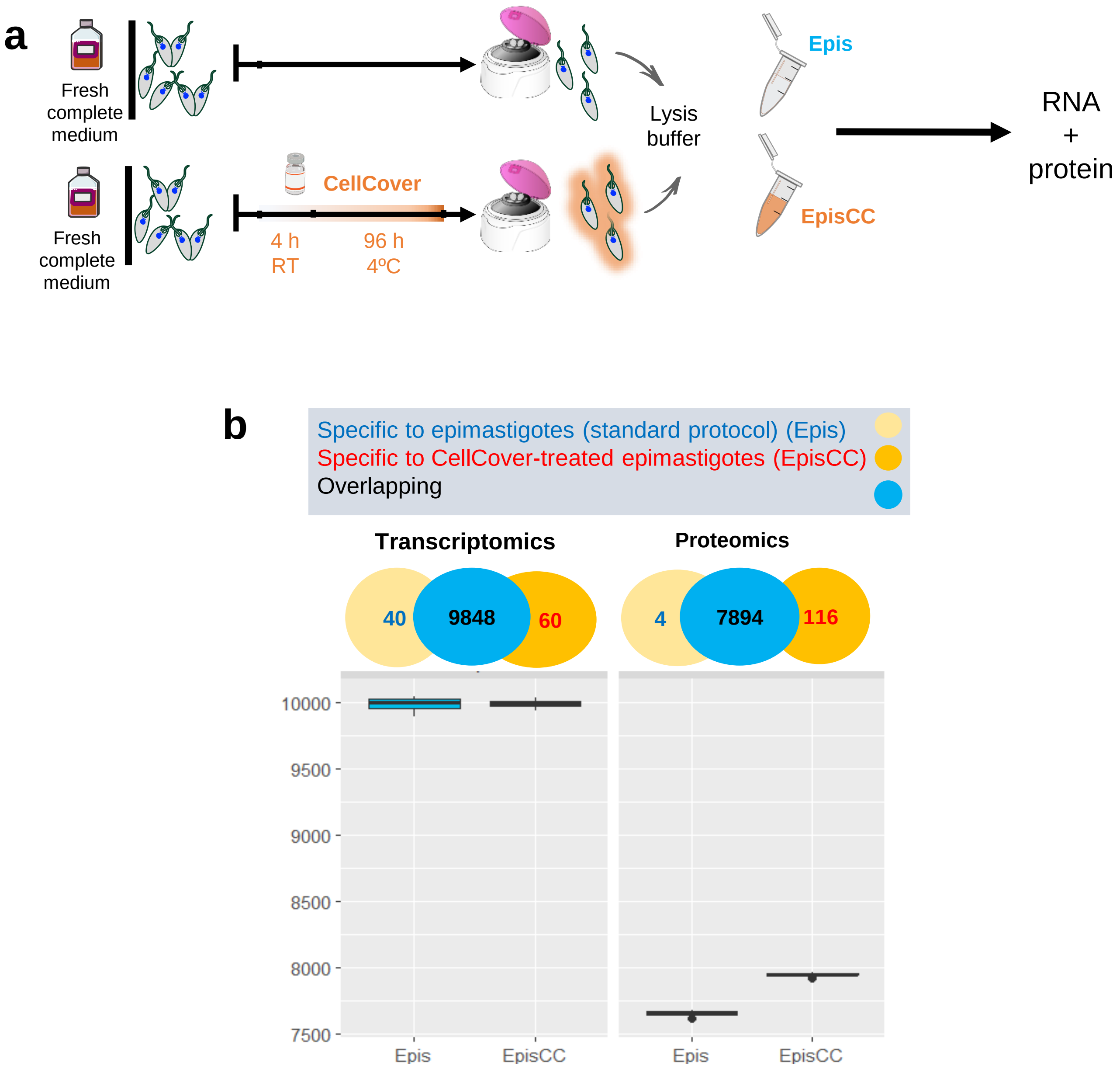


Extended Data Fig. 3

**a**

| | MCM2 | MCM3 | MCM4 | MCM5 | MCM6 | MCM7 |
|---|---|---|---|---|---|---|
| Epi RNA | 1.00 | 1.00 | 1.00 | 1.00 | 1.00 | 1.00 |
| QEpi RNA | 0.24 | 0.24 | 0.26 | 0.20 | 0.32 | 0.32 |

**b**

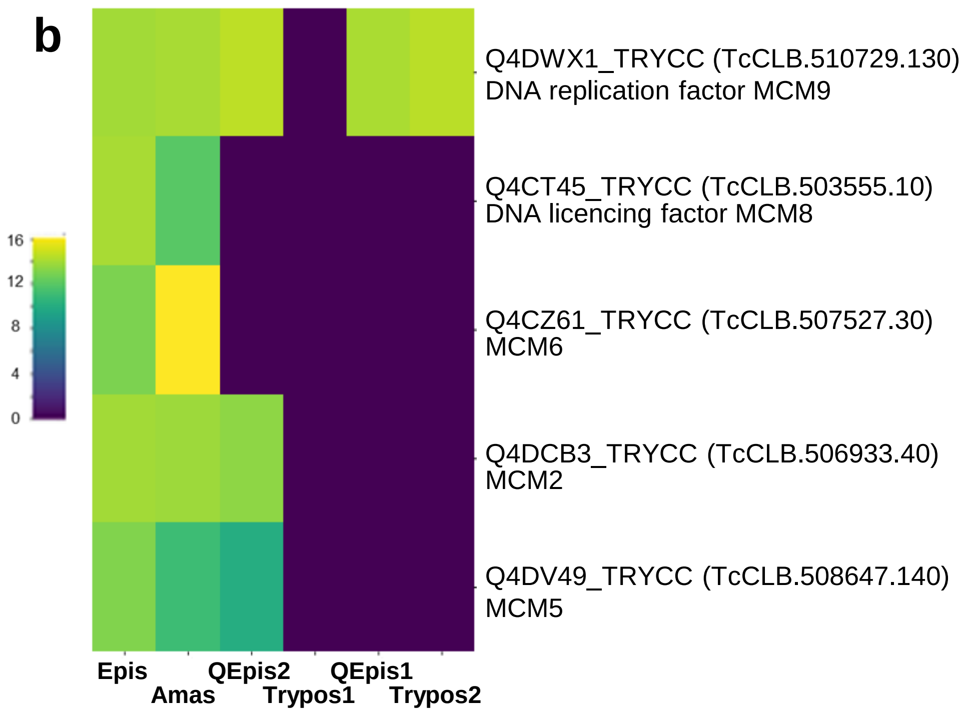


Extended Data Fig. 4

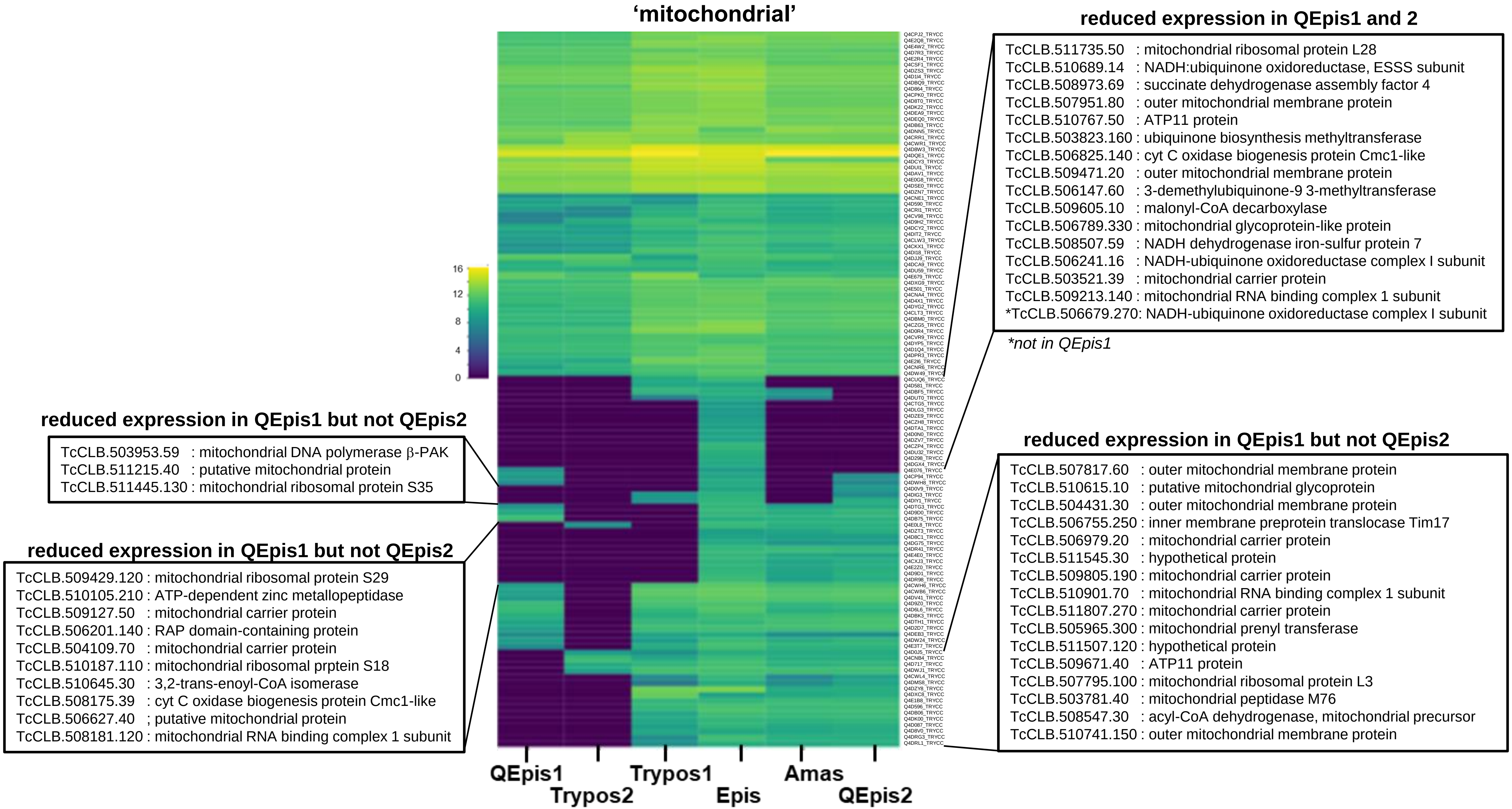


Extended Data Fig. 5

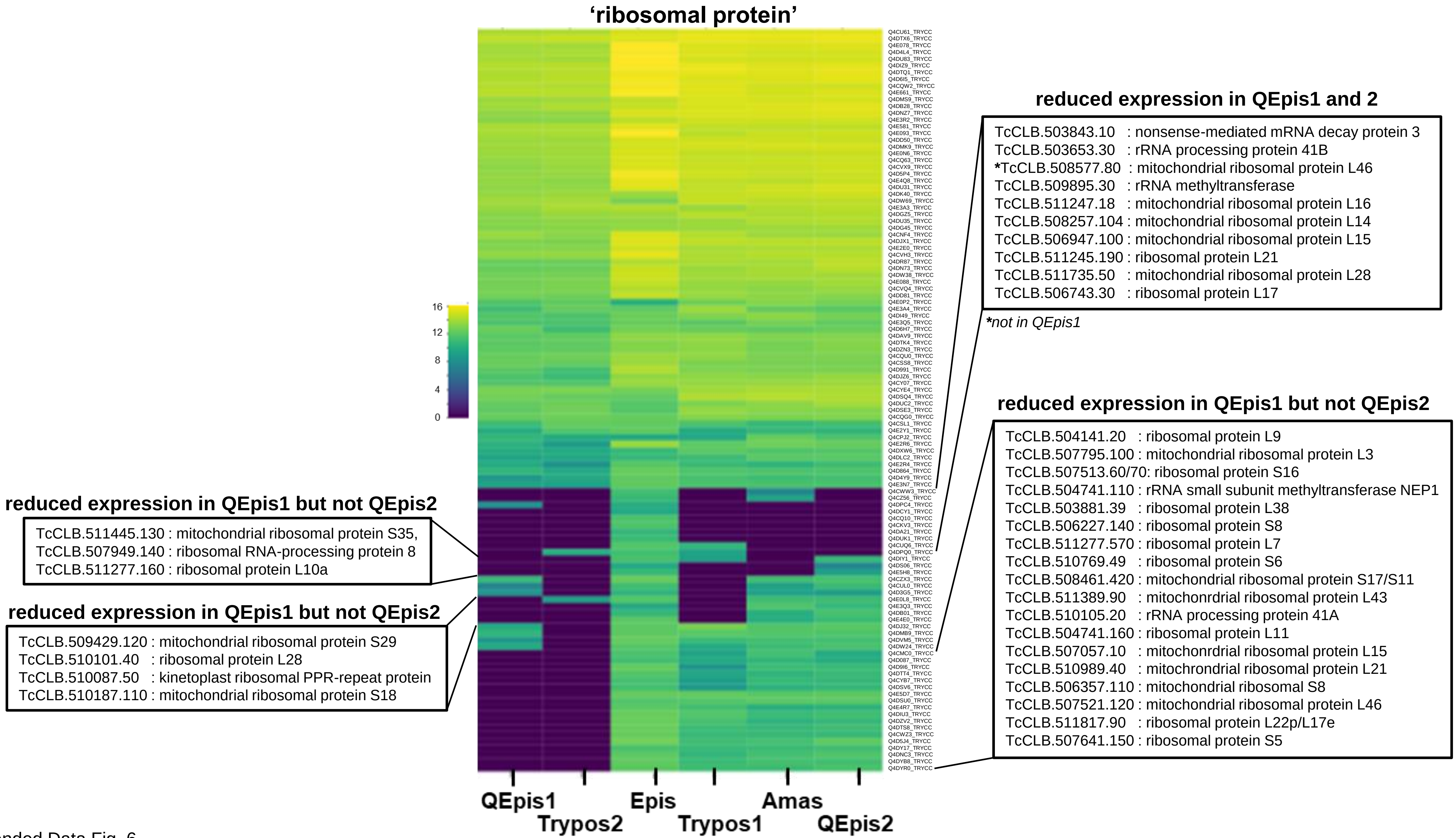


Extended Data Fig. 6

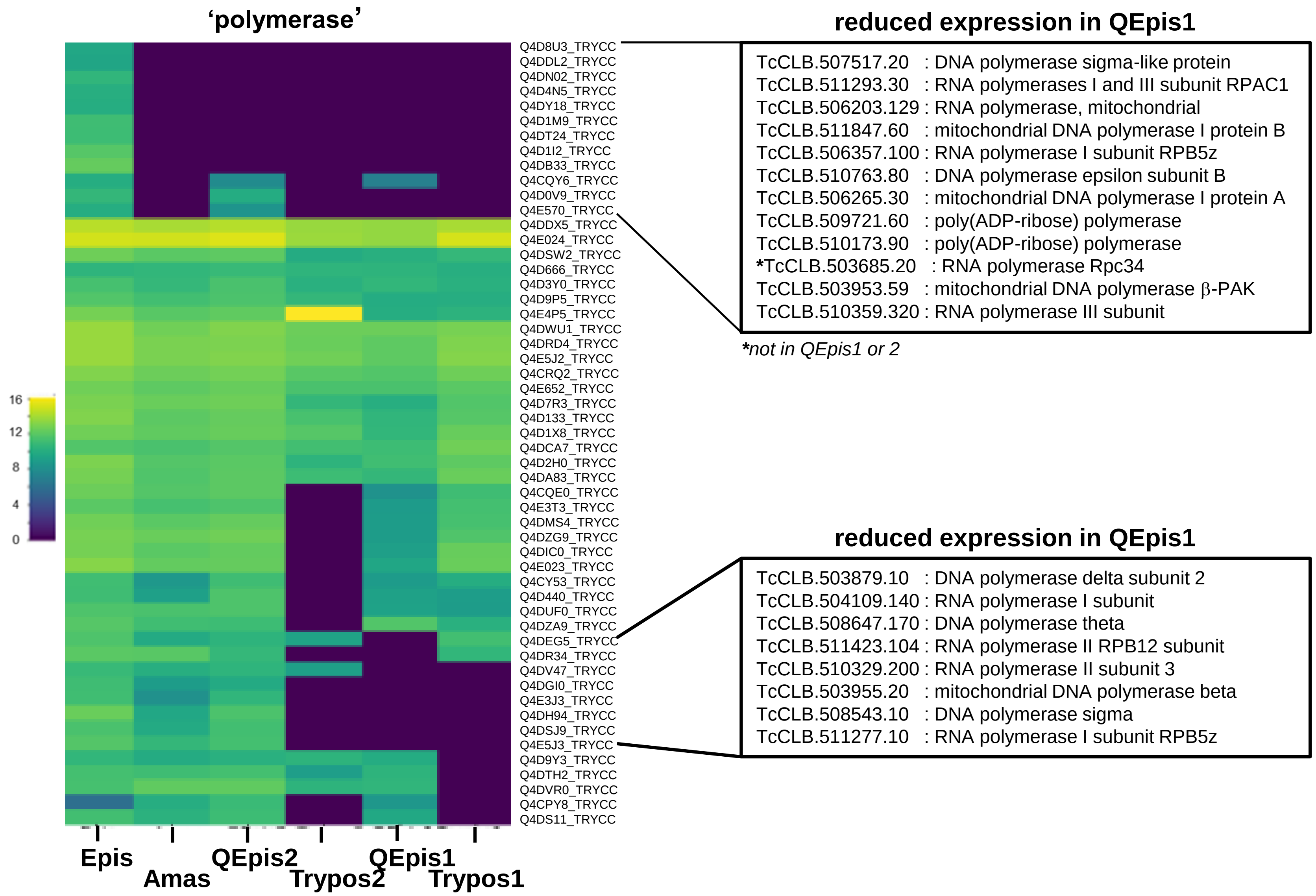


Extended Data Fig. 7

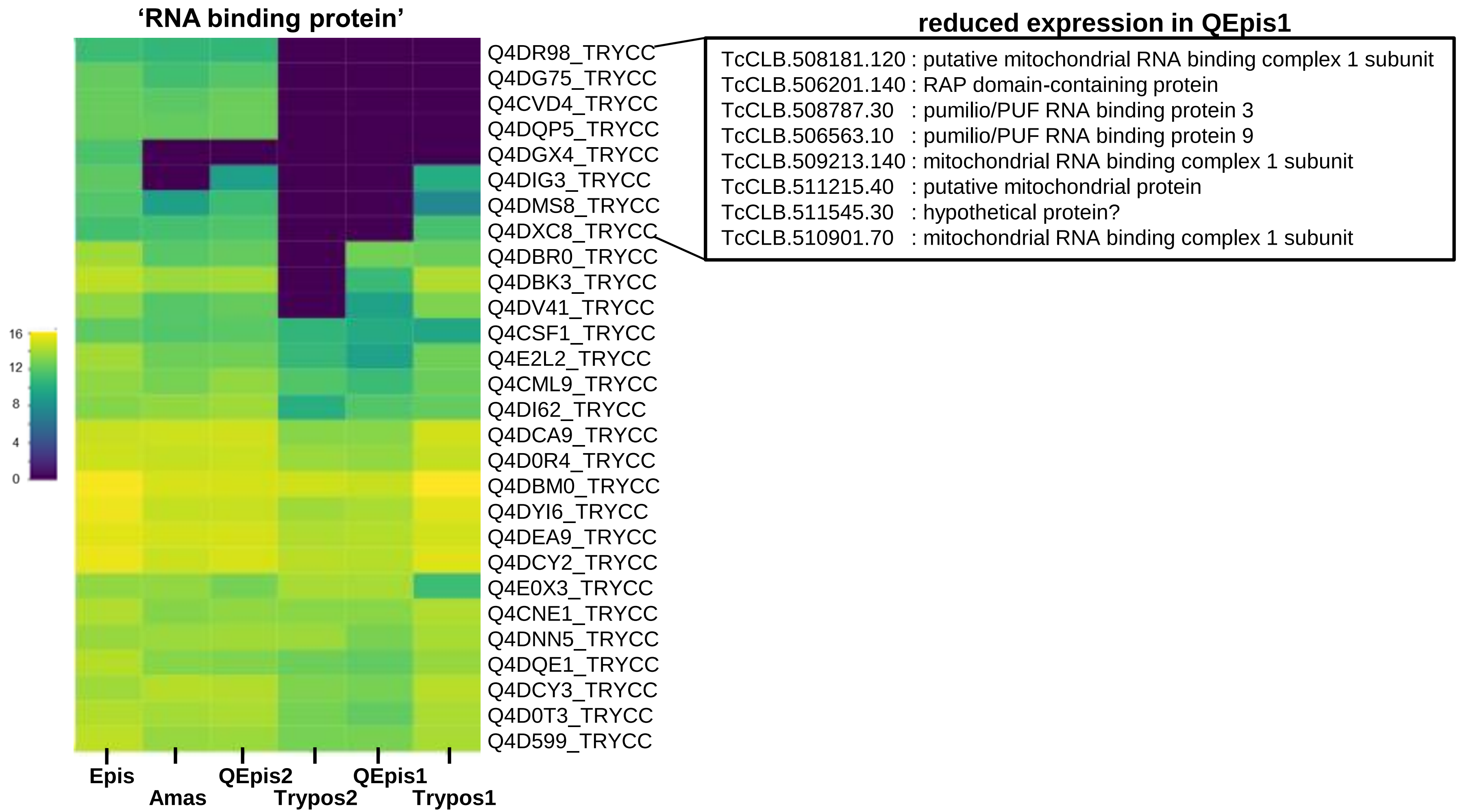


Extended data Fig. 8
