## Supplementary Tables 1-3 for "Transcriptomic and proteomic analysis of quiescent epimastigotes as a resource for investigating *Trypanosoma cruzi* persistence"

**Supplementary Table 1. Genes up-regulated (brown shading) and down-regulated (blue shading) in QEpis relative to Epis, as inferred from the transcriptome data (log2FC >2 and adj. p-value <0.05)**

| accession number | log2FC | putative gene function |
|---|---|---|
| Tc00.1047053511805.10 | 5.06 | mannosyl-oligosaccharide glucosidase, glucosidase I, processing A-glucosidase I |
| Tc00.1047053511357.5 | 5.04 | glycerol uptake protein |
| Tc00.1047053506979.5 | 4.51 | hypothetical protein, conserved (pseudogene) |
| Tc00.1047053511001.18 | 4.32 | 40S ribosomal protein S3A |
| Tc00.1047053511369.30 | 4.26 | elongation factor 1-alpha (ef-1-alpha) |
| Tc00.1047053503859.60 | 4.15 | hypothetical protein, conserved (pseudogene) |
| Tc00.1047053506907.40 | 4.03 | hypothetical protein |
| Tc00.1047053508653.20 | 4.01 | ribosomal RNA small subunit, 5' partial |
| Tc00.1047053510157.5 | 3.92 | trans-sialidase |
| Tc00.1047053511705.5 | 3.82 | trans-sialidase (pseudogene) |
| Tc00.1047053507957.304 | 3.76 | surface protease GP63 (pseudogene) |
| Tc00.1047053506565.8 | 3.05 | RNA-binding protein, orthologue of TbRNP7 |
| Tc00.1047053507145.98 | 3.05 | hypothetical protein |
| Tc00.1047053508157.40 | 3.01 | hypothetical protein |
| Tc00.1047053460061.10 | 3.00 | hypothetical protein |
| Tc00.1047053421173.14 | 2.98 | trans-sialidase |
| Tc00.1047053508087.10 | 2.94 | hypothetical protein |
| Tc00.1047053511121.20 | 2.93 | hypothetical protein |
| Tc00.1047053424795.10 | 2.86 | hypothetical protein, conserved |
| Tc00.1047053507957.310 | 2.85 | hypothetical protein |
| Tc00.1047053504239.90 | 2.83 | hypothetical protein, conserved |
| Tc00.1047053508321.11 | 2.81 | histone H2A |
| Tc00.1047053504153.50 | 2.80 | ribosomal RNA large subunit alpha, 5' & 3' partial |
| Tc00.1047053508687.30 | 2.74 | mucin-associated surface protein (MASP, pseudogene) |
| Tc00.1047053511603.434 | 2.72 | surface protease GP63 |
| Tc00.1047053506343.51 | 2.69 | retrotransposon hot spot protein (RHS, pseudogene), |
| Tc00.1047053508165.250 | 2.69 | hypothetical protein, conserved |
| Tc00.1047053506499.130 | 2.66 | hypothetical protein |
| Tc00.1047053511127.74 | 2.65 | hypothetical protein, conserved |
| Tc00.1047053508229.50 | 2.62 | hypothetical protein |
| Tc00.1047053432621.9 | 2.62 | dispersed gene family protein 1 (DGF-1) |
| Tc00.1047053509513.5 | 2.62 | hypothetical protein, conserved (pseudogene) |
| Tc00.1047053506579.129 | 2.60 | hypothetical protein, conserved |
| Tc00.1047053508143.110 | 2.57 | hypothetical protein |
| Tc00.1047053506267.10 | 2.51 | mucin-associated surface protein (MASP) |
| Tc00.1047053506857.120 | 2.48 | hypothetical protein, conserved |
| Tc00.1047053479517.50 | 2.46 | surface protease GP63 |
| Tc00.1047053503599.60 | 2.45 | hypothetical protein, conserved |
| Tc00.1047053507237.350 | 2.42 | mucin TcMUCII |

| | | |
|---|---|---|
| Tc00.1047053422207.10 | 2.41 | UDP-Gal or UDP-GlcNAc-dependent glycosyltransferase |
| Tc00.1047053509631.11 | 2.40 | hypothetical protein |
| Tc00.1047053508777.210 | 2.39 | hypothetical protein |
| Tc00.1047053506431.5 | 2.30 | trans-sialidase, |
| Tc00.1047053510749.99 | 2.27 | aminopeptidase P |
| Tc00.1047053510275.21 | 2.24 | surface protease GP63 (pseudogene) |
| Tc00.1047053508097.10 | 2.23 | hypothetical protein |
| Tc00.1047053511173.340 | 2.22 | mucin TcMUCII (pseudogene) |
| Tc00.1047053506973.140 | 2.16 | hypothetical protein |
| Tc00.1047053507143.90 | 2.16 | hypothetical protein |
| Tc00.1047053510275.180 | 2.13 | elongation factor 1-gamma (EF-1-gamma) |
| Tc00.1047053510105.131 | 2.13 | C/D small nucleolar RNA (snoRNA), TB8C2C1 |
| Tc00.1047053506965.120 | 2.11 | hypothetical protein |
| Tc00.1047053506979.30 | 2.11 | hypothetical protein |
| Tc00.1047053508107.20 | 2.10 | trans-sialidase (pseudogene) |
| Tc00.1047053510269.40 | 2.08 | hypothetical protein |
| Tc00.1047053507755.10 | 2.06 | hypothetical protein |
| Tc00.1047053504039.110 | 2.06 | hypothetical protein |
| Tc00.1047053506763.370 | 2.06 | hypothetical protein |
| Tc00.1047053508209.120 | 2.06 | 10 kDa heat shock protein |
| Tc00.1047053506969.6 | 2.01 | trans-sialidase (pseudogene) |
| Tc00.1047053508165.360 | -2.01 | surface protease GP63 (pseudogene) |
| Tc00.1047053507547.18 | -2.01 | mucin TcMUCI |
| Tc00.1047053506811.60 | -2.01 | hypothetical protein |
| Tc00.1047053511613.166 | -2.01 | surface protease GP63 (pseudogene) |
| Tc00.1047053510181.140 | -2.01 | hypothetical protein, conserved |
| Tc00.1047053510021.30 | -2.02 | mucin TcMUCII |
| Tc00.1047053506499.90 | -2.02 | 90 kDa surface protein, serine-alanine-and proline-rich protein |
| Tc00.1047053506965.190 | -2.02 | mucin TcMUCII |
| Tc00.1047053506501.120 | -2.02 | mucin TcMUCII |
| Tc00.1047053506285.50 | -2.03 | mucin TcMUCII |
| Tc00.1047053510959.15 | -2.03 | elongation factor 1-gamma (EF-1-gamma, pseudogene) |
| Tc00.1047053507521.115 | -2.03 | hypothetical protein, conserved |
| Tc00.1047053510013.290 | -2.03 | mucin-associated surface protein (MASP) |
| Tc00.1047053510213.39 | -2.03 | mucin-associated surface protein (MASP) |
| Tc00.1047053507987.20 | -2.04 | hypothetical protein |
| Tc00.1047053510289.54 | -2.04 | hypothetical protein, conserved |
| Tc00.1047053503619.25 | -2.04 | hypothetical protein, conserved |
| Tc00.1047053506755.240 | -2.04 | hypothetical protein, conserved |
| Tc00.1047053511603.190 | -2.05 | mucin-associated surface protein (MASP) |
| Tc00.1047053511797.150 | -2.05 | hypothetical protein |
| Tc00.1047053504103.60 | -2.05 | hypothetical protein, conserved |
| Tc00.1047053504239.380 | -2.05 | hypothetical protein |
| Tc00.1047053511553.45 | -2.06 | surface protease GP63 (pseudogene) |
| Tc00.1047053510561.60 | -2.06 | hypothetical protein |
| Tc00.1047053508165.110 | -2.06 | mucin-associated surface protein (MASP) |

| | | |
|---|---|---|
| Tc00.1047053509925.20 | -2.06 | mucin TcMUCII |
| Tc00.1047053506865.4 | -2.06 | hypothetical protein, conserved |
| Tc00.1047053510375.40 | -2.06 | mucin-associated surface protein (MASP) |
| Tc00.1047053509197.39 | -2.07 | cation transporter |
| Tc00.1047053504239.340 | -2.07 | mucin TcMUCII |
| Tc00.1047053507953.210 | -2.07 | hypothetical protein |
| Tc00.1047053508977.31 | -2.08 | elongation factor 1-gamma (EF-1-gamma, pseudogene) |
| Tc00.1047053510373.90 | -2.08 | mucin-associated surface protein (MASP) |
| Tc00.1047053511603.80 | -2.08 | mucin-associated surface protein (MASP) |
| Tc00.1047053505025.100 | -2.08 | mucin TcMUCII (pseudogene) |
| Tc00.1047053504239.270 | -2.08 | hypothetical protein |
| Tc00.1047053506767.324 | -2.08 | mucin TcMUCII |
| Tc00.1047053511685.50 | -2.08 | hypothetical protein, conserved |
| Tc00.1047053511605.55 | -2.08 | hypothetical protein |
| Tc00.1047053506717.240 | -2.08 | hypothetical protein, conserved |
| Tc00.1047053507163.30 | -2.09 | 90 kDa surface protein, serine-alanine-and proline-rich protein |
| Tc00.1047053508047.80 | -2.09 | mucin TcMUCII (pseudogene) |
| Tc00.1047053508163.105 | -2.09 | elongation factor 1-gamma (EF-1-gamma, pseudogene) |
| Tc00.1047053509443.39 | -2.09 | hypothetical protein, conserved |
| Tc00.1047053508081.80 | -2.09 | hypothetical protein |
| Tc00.1047053506667.120 | -2.09 | mucin-associated surface protein (MASP) |
| Tc00.1047053510275.355 | -2.10 | hypothetical protein |
| Tc00.1047053508283.60 | -2.10 | retrotransposon hot spot protein (RHS, pseudogene) |
| Tc00.1047053503503.40 | -2.10 | mucin-associated surface protein (MASP) |
| Tc00.1047053507237.110 | -2.10 | elongation factor 1-gamma (EF-1-gamma) |
| Tc00.1047053506599.224 | -2.10 | mucin TcMUCII |
| Tc00.1047053506267.101 | -2.10 | surface protease GP63 (pseudogene) |
| Tc00.1047053503971.30 | -2.10 | hypothetical protein, conserved |
| Tc00.1047053506599.360 | -2.11 | mucin-associated surface protein (MASP, pseudogene) |
| Tc00.1047053507959.300 | -2.11 | mucin-associated surface protein (MASP) |
| Tc00.1047053511603.60 | -2.11 | mucin TcMUCII |
| Tc00.1047053510603.60 | -2.11 | protein tyrosine phosphatase-like protein |
| Tc00.1047053511739.30 | -2.12 | hypothetical protein, conserved |
| Tc00.1047053510175.125 | -2.12 | hypothetical protein |
| Tc00.1047053507009.100 | -2.12 | cyclophilin, PPIase, rotamase, peptidyl-prolyl cis-trans isomerase |
| Tc00.1047053506759.210 | -2.13 | mucin TcMUCII |
| Tc00.1047053479721.10 | -2.13 | hypothetical protein, conserved (pseudogene) |
| Tc00.1047053510013.70 | -2.13 | mucin-associated surface protein (MASP) |
| Tc00.1047053506599.430 | -2.13 | mucin TcMUCII |
| Tc00.1047053506667.30 | -2.14 | 90 kDa surface protein |
| Tc00.1047053504507.20 | -2.14 | DNA replication licensing factor, minichromosome maintenance protein-like protein |
| Tc00.1047053508307.40 | -2.15 | hypothetical protein, conserved |
| Tc00.1047053510375.30 | -2.16 | hypothetical protein, conserved (pseudogene) |
| Tc00.1047053508097.60 | -2.16 | mucin-associated surface protein (MASP) |
| Tc00.1047053508439.40 | -2.16 | hypothetical protein |
| Tc00.1047053504039.200 | -2.17 | mucin-associated surface protein (MASP) |

| | | |
|---|---|---|
| Tc00.1047053506289.150 | -2.17 | mucin-associated surface protein (MASP) |
| Tc00.1047053510013.180 | -2.18 | hypothetical protein |
| Tc00.1047053509871.21 | -2.18 | N-acetyltransferase complex ARD1 subunit (pseudogene) |
| Tc00.1047053503759.30 | -2.18 | trans-sialidase |
| Tc00.1047053506971.10 | -2.19 | mucin-associated surface protein (MASP) |
| Tc00.1047053510143.17 | -2.19 | hypothetical protein, conserved |
| Tc00.1047053506291.20 | -2.19 | hypothetical protein |
| Tc00.1047053504239.40 | -2.19 | surface protease GP63 (pseudogene) |
| Tc00.1047053511173.370 | -2.19 | trans-sialidase |
| Tc00.1047053506267.19 | -2.19 | hypothetical protein |
| Tc00.1047053510157.10 | -2.20 | mucin-associated surface protein (MASP), |
| Tc00.1047053504239.280 | -2.20 | 90 kDa surface protein, serine-alanine-and proline-rich protein |
| Tc00.1047053504039.120 | -2.20 | mucin TcMUCII |
| Tc00.1047053508157.20 | -2.21 | mucin-associated surface protein (MASP) |
| Tc00.1047053510371.110 | -2.21 | mucin TcMUCII |
| Tc00.1047053508109.30 | -2.22 | mucin-associated surface protein (MASP) |
| Tc00.1047053511401.111 | -2.23 | elongation factor 1-gamma (EF-1-gamma, pseudogene) |
| Tc00.1047053510371.100 | -2.23 | mucin-associated surface protein (MASP) |
| Tc00.1047053508681.11 | -2.23 | trans-sialidase (pseudogene) |
| Tc00.1047053511613.80 | -2.23 | mucin-associated surface protein (MASP) |
| Tc00.1047053505945.20 | -2.24 | ribonuclease mar1 |
| Tc00.1047053507237.80 | -2.24 | mucin-associated surface protein (MASP) |
| Tc00.1047053403153.10 | -2.24 | receptor-type adenylate cyclase (pseudogene) |
| Tc00.1047053506501.150 | -2.24 | surface protease GP63 (pseudogene) |
| Tc00.1047053508109.14 | -2.24 | mucin TcMUCII |
| Tc00.1047053508379.29 | -2.24 | hypothetical protein, conserved |
| Tc00.1047053508047.40 | -2.25 | mucin TcMUCII (pseudogene) |
| Tc00.1047053506769.60 | -2.25 | mucin-associated surface protein (MASP, pseudogene) |
| Tc00.1047053510373.110 | -2.25 | hypothetical protein |
| Tc00.1047053507163.20 | -2.26 | hypothetical protein |
| Tc00.1047053509769.10 | -2.26 | hypothetical protein, conserved |
| Tc00.1047053511679.10 | -2.27 | mucin TcSMUGS |
| Tc00.1047053506597.33 | -2.27 | hypothetical protein |
| Tc00.1047053510375.5 | -2.27 | surface protease GP63 (pseudogene) |
| Tc00.1047053506885.410 | -2.28 | hypothetical protein, conserved |
| Tc00.1047053508165.50 | -2.28 | mucin-associated surface protein (MASP) |
| Tc00.1047053509127.59 | -2.28 | hypothetical protein, conserved |
| Tc00.1047053511529.20 | -2.29 | hypothetical protein, conserved |
| Tc00.1047053511613.50 | -2.30 | mucin TcMUCII |
| Tc00.1047053503531.40 | -2.30 | hypothetical protein, conserved |
| Tc00.1047053510525.80 | -2.31 | histone H2A |
| Tc00.1047053506409.190 | -2.31 | mucin-associated surface protein (MASP, pseudogene) |
| Tc00.1047053510021.90 | -2.31 | hypothetical protein, conserved (pseudogene) |
| Tc00.1047053511171.25 | -2.32 | hypothetical protein, conserved (pseudogene) |
| Tc00.1047053507237.170 | -2.32 | mucin-associated surface protein (MASP) |
| Tc00.1047053508307.100 | -2.33 | hypothetical protein, conserved |

| | | |
|---|---|---|
| Tc00.1047053507611.210 | -2.33 | hypothetical protein |
| Tc00.1047053503937.40 | -2.33 | trans-sialidase (pseudogene) |
| Tc00.1047053511603.390 | -2.34 | mucin TcMUCII |
| Tc00.1047053503977.15 | -2.34 | surface protease GP63 (pseudogene) |
| Tc00.1047053503909.110 | -2.34 | intraflagellar transport (IFT) protein |
| Tc00.1047053507237.100 | -2.34 | mucin-associated surface protein (MASP) |
| Tc00.1047053511729.70 | -2.34 | cationic amino acid transporter |
| Tc00.1047053508165.34 | -2.34 | mucin TcMUC (pseudogene) |
| Tc00.1047053510021.100 | -2.34 | mucin TcMUCII |
| Tc00.1047053508207.54 | -2.35 | hypothetical protein, conserved |
| Tc00.1047053506269.59 | -2.35 | hypothetical protein (pseudogene) |
| Tc00.1047053507011.150 | -2.35 | hypothetical protein, conserved |
| Tc00.1047053511877.15 | -2.36 | elongation factor 1-gamma (EF-1-gamma, pseudogene) |
| Tc00.1047053507699.149 | -2.36 | mucin TcMUCII |
| Tc00.1047053506965.180 | -2.36 | mucin-associated surface protein (MASP) |
| Tc00.1047053506767.400 | -2.36 | mucin-associated surface protein (MASP) |
| Tc00.1047053510143.129 | -2.36 | hypothetical protein, conserved |
| Tc00.1047053511213.15 | -2.37 | mucin-associated surface protein (MASP) |
| Tc00.1047053422319.10 | -2.37 | hypothetical protein |
| Tc00.1047053507959.270 | -2.38 | mucin TcMUCII |
| Tc00.1047053507993.89 | -2.38 | hypothetical protein, conserved |
| Tc00.1047053507641.240 | -2.38 | hypothetical protein |
| Tc00.1047053506601.31 | -2.39 | surface protease GP63 (pseudogene) |
| Tc00.1047053511625.90 | -2.39 | mucin-associated surface protein (MASP) |
| Tc00.1047053508685.9 | -2.39 | dispersed gene family protein 1 (DGF-1) |
| Tc00.1047053507747.110 | -2.39 | mucin TcMUCII |
| Tc00.1047053511433.20 | -2.39 | hypothetical protein, conserved |
| Tc00.1047053505025.10 | -2.40 | mucin-associated surface protein (MASP) |
| Tc00.1047053509631.30 | -2.41 | mucin TcMUCII (pseudogene) |
| Tc00.1047053506181.97 | -2.42 | hypothetical protein, conserved |
| Tc00.1047053507641.260 | -2.42 | HSP60, mitochondrial precursor (pseudogene) groELprotein, frameshift, |
| Tc00.1047053508647.174 | -2.42 | hypothetical protein, conserved |
| Tc00.1047053508227.51 | -2.42 | surface protease GP63 (pseudogene) |
| Tc00.1047053511603.200 | -2.42 | mucin TcMUCII |
| Tc00.1047053506767.196 | -2.42 | surface protease GP63 (pseudogene) |
| Tc00.1047053507957.90 | -2.42 | mucin TcMUCII |
| Tc00.1047053510025.165 | -2.43 | hypothetical protein, conserved (pseudogene) |
| Tc00.1047053506667.130 | -2.43 | surface protease GP63 (pseudogene) |
| Tc00.1047053510261.31 | -2.43 | mucin-associated surface protein (MASP, pseudogene) |
| Tc00.1047053410589.19 | -2.43 | hypothetical protein, conserved |
| Tc00.1047053511603.110 | -2.43 | mucin TcMUCII |
| Tc00.1047053507237.180 | -2.44 | hypothetical protein |
| Tc00.1047053510561.14 | -2.45 | hypothetical protein |
| Tc00.1047053509103.4 | -2.45 | protein kinase |
| Tc00.1047053508165.400 | -2.46 | mucin TcMUCII |
| Tc00.1047053506925.370 | -2.47 | MP44 (pseudogene) |

| | | |
|---|---|---|
| Tc00.1047053506737.60 | -2.47 | hypothetical protein, conserved |
| Tc00.1047053506113.55 | -2.47 | N-acetyltransferase complex ARD1 subunit (pseudogene) |
| Tc00.1047053510105.322 | -2.47 | elongation factor 1-gamma (EF-1-gamma, pseudogene) |
| Tc00.1047053510089.5 | -2.47 | mucin TcMUCII (pseudogene) |
| Tc00.1047053509827.4 | -2.48 | retrotransposon hot spot (RHS) protein |
| Tc00.1047053510021.80 | -2.49 | 90 kDa surface protein |
| Tc00.1047053508081.10 | -2.49 | mucin-associated surface protein (MASP) |
| Tc00.1047053504039.90 | -2.50 | mucin-associated surface protein (MASP, pseudogene) |
| Tc00.1047053506501.160 | -2.50 | mucin-associated surface protein (MASP, pseudogene) |
| Tc00.1047053508151.20 | -2.50 | mucin-associated surface protein (MASP) |
| Tc00.1047053508147.70 | -2.50 | trans-sialidase (pseudogene) |
| Tc00.1047053507485.80 | -2.50 | hypothetical protein, conserved |
| Tc00.1047053511621.200 | -2.52 | ribosomal protein L11 |
| Tc00.1047053507041.85 | -2.52 | hypothetical protein, conserved (pseudogene) |
| Tc00.1047053506529.425 | -2.52 | hypothetical protein |
| Tc00.1047053510055.20 | -2.52 | trans-sialidase |
| Tc00.1047053507699.180 | -2.53 | mucin-associated surface protein (MASP, pseudogene) |
| Tc00.1047053508165.140 | -2.54 | mucin TcMUCII (pseudogene) |
| Tc00.1047053508229.20 | -2.55 | hypothetical protein |
| Tc00.1047053506367.10 | -2.55 | isovaleryl-coA dehydrogenase |
| Tc00.1047053506501.280 | -2.55 | mucin TcMUCII |
| Tc00.1047053506599.170 | -2.55 | mucin-associated surface protein (MASP) |
| Tc00.1047053506765.74 | -2.55 | mucin-associated surface protein (MASP) |
| Tc00.1047053503717.31 | -2.55 | mucin TcMUCII |
| Tc00.1047053511211.80 | -2.56 | hypothetical protein, conserved |
| Tc00.1047053507957.60 | -2.56 | mucin-associated surface protein (MASP, pseudogene) |
| Tc00.1047053503751.20 | -2.56 | hypothetical protein |
| Tc00.1047053505807.209 | -2.57 | hypothetical protein, conserved |
| Tc00.1047053510261.41 | -2.57 | mucin TcMUC (pseudogene) |
| Tc00.1047053503503.15 | -2.58 | hypothetical protein (pseudogene) |
| Tc00.1047053506599.130 | -2.58 | mucin TcMUCII (pseudogene) |
| Tc00.1047053510191.20 | -2.59 | mucin TcMUCII |
| Tc00.1047053510189.30 | -2.59 | surface protease GP63 (pseudogene) |
| Tc00.1047053510279.40 | -2.59 | 90 kDa surface protein, serine-alanine-and proline-rich protein |
| Tc00.1047053510107.41 | -2.60 | elongation factor 1-gamma (EF-1-gamma, pseudogene) |
| Tc00.1047053504125.40 | -2.61 | ATPase subunit 9 |
| Tc00.1047053506973.30 | -2.61 | mucin-associated surface protein (MASP) |
| Tc00.1047053509699.85 | -2.61 | surface protease GP63 (pseudogene) |
| Tc00.1047053506743.69 | -2.61 | hypothetical protein, conserved |
| Tc00.1047053511553.5 | -2.62 | trans-sialidase (pseudogene) |
| Tc00.1047053511611.40 | -2.63 | hypothetical protein |
| Tc00.1047053507953.40 | -2.64 | mucin-associated surface protein (MASP) |
| Tc00.1047053508375.60 | -2.64 | succinate dehydrogenase subunit |
| Tc00.1047053511471.10 | -2.65 | hypothetical protein, conserved (pseudogene) |
| Tc00.1047053507955.20 | -2.65 | mucin-associated surface protein (MASP) |
| Tc00.1047053511585.180 | -2.65 | hypothetical protein, conserved |

| | | |
|---|---|---|
| Tc00.1047053505025.60 | -2.65 | 90 kDa surface protein |
| Tc00.1047053510431.290 | -2.65 | hypothetical protein |
| Tc00.1047053511735.40 | -2.65 | hypothetical protein, conserved |
| Tc00.1047053469435.4 | -2.67 | ribosomal protein L14 |
| Tc00.1047053511553.90 | -2.67 | mucin TcMUCII |
| Tc00.1047053510013.240 | -2.68 | mucin TcMUCII |
| Tc00.1047053510279.180 | -2.69 | mucin TcMUCII (pseudogene) |
| Tc00.1047053511121.30 | -2.69 | hypothetical protein |
| Tc00.1047053508207.120 | -2.70 | hypothetical protein, conserved |
| Tc00.1047053507237.24 | -2.71 | mucin TcMUCII (pseudogene) |
| Tc00.1047053506267.70 | -2.72 | mucin-associated surface protein (MASP) |
| Tc00.1047053507959.224 | -2.72 | surface protease GP63 (pseudogene) |
| Tc00.1047053508721.4 | -2.72 | hypothetical protein, conserved |
| Tc00.1047053511211.30 | -2.73 | hypothetical protein |
| Tc00.1047053506269.79 | -2.74 | mucin-associated surface protein (MASP, pseudogene) |
| Tc00.1047053506529.314 | -2.74 | hypothetical protein, conserved |
| Tc00.1047053503689.34 | -2.74 | hypothetical protein, conserved |
| Tc00.1047053511173.440 | -2.74 | trans-sialidase |
| Tc00.1047053508977.111 | -2.75 | elongation factor 1-gamma (EF-1-gamma, pseudogene) |
| Tc00.1047053508099.20 | -2.75 | serine-alanine-and proline-rich protein |
| Tc00.1047053510021.160 | -2.75 | 90 kDa surface protein |
| Tc00.1047053510371.70 | -2.76 | mucin-associated surface protein (MASP) |
| Tc00.1047053506767.119 | -2.76 | surface protease GP63 (pseudogene) |
| Tc00.1047053503835.20 | -2.76 | hypothetical protein, conserved |
| Tc00.1047053510101.390 | -2.76 | hypothetical protein, conserved |
| Tc00.1047053503501.70 | -2.76 | mucin-associated surface protein (MASP) |
| Tc00.1047053507041.50 | -2.77 | hypothetical protein, conserved |
| Tc00.1047053511733.100 | -2.77 | hypothetical protein, conserved |
| Tc00.1047053510557.76 | -2.78 | hypothetical protein |
| Tc00.1047053510999.39 | -2.79 | 40S ribosomal protein S3A |
| Tc00.1047053510281.5 | -2.80 | mucin TcMUCII (pseudogene) |
| Tc00.1047053503501.60 | -2.81 | hypothetical protein, conserved (pseudogene) |
| Tc00.1047053507047.180 | -2.81 | hypothetical protein, conserved |
| Tc00.1047053510191.80 | -2.81 | mucin TcMUCII |
| Tc00.1047053508099.30 | -2.82 | hypothetical protein |
| Tc00.1047053511589.260 | -2.84 | glycyl-tRNA synthetase |
| Tc00.1047053503601.20 | -2.85 | hypothetical protein |
| Tc00.1047053510261.21 | -2.85 | hypothetical protein |
| Tc00.1047053509079.20 | -2.85 | mucin-associated surface protein (MASP) |
| Tc00.1047053508163.40 | -2.86 | mucin TcMUCII (pseudogene) |
| Tc00.1047053511599.30 | -2.86 | 90 kDa surface protein |
| Tc00.1047053510377.220 | -2.87 | mucin-associated surface protein (MASP, pseudogene) |
| Tc00.1047053508097.81 | -2.90 | mucin TcMUCII |
| Tc00.1047053506769.55 | -2.92 | hypothetical protein |
| Tc00.1047053504239.420 | -2.92 | mucin-associated surface protein (MASP) |
| Tc00.1047053511607.110 | -2.93 | mucin-associated surface protein (MASP) |

| | | |
|---|---|---|
| Tc00.1047053506599.91 | -2.93 | surface protease GP63 (pseudogene) |
| Tc00.1047053509195.50 | -2.94 | mucin-associated surface protein (MASP) |
| Tc00.1047053510279.130 | -2.96 | mucin TcMUCII |
| Tc00.1047053510371.30 | -2.96 | hypothetical protein, conserved (pseudogene) |
| Tc00.1047053507953.180 | -2.96 | mucin TcMUCII |
| Tc00.1047053511597.40 | -2.97 | mucin-associated surface protein (MASP) |
| Tc00.1047053511599.110 | -2.98 | mucin TcMUCII |
| Tc00.1047053506829.5 | -2.99 | hypothetical protein |
| Tc00.1047053508389.130 | -2.99 | mucin-associated surface protein (MASP) |
| Tc00.1047053506609.80 | -2.99 | mucin TcMUCII |
| Tc00.1047053504163.50 | -3.01 | fructose-bisphosphate aldolase, glycosomal |
| Tc00.1047053511437.20 | -3.02 | expression site-associated gene (ESAG-like) protein |
| Tc00.1047053510317.34 | -3.06 | poly(A) polymerase |
| Tc00.1047053510371.61 | -3.07 | surface protease GP63 (pseudogene) |
| Tc00.1047053506501.350 | -3.09 | mucin-associated surface protein (MASP) |
| Tc00.1047053506499.100 | -3.09 | hypothetical protein |
| Tc00.1047053510191.10 | -3.10 | mucin-associated surface protein (MASP) |
| Tc00.1047053506545.5 | -3.11 | mucin TcMUCI |
| Tc00.1047053510377.120 | -3.11 | hypothetical protein |
| Tc00.1047053506363.159 | -3.12 | Separase |
| Tc00.1047053506971.60 | -3.12 | mucin-associated surface protein (MASP) |
| Tc00.1047053506501.290 | -3.15 | mucin-associated surface protein (MASP) |
| Tc00.1047053510377.310 | -3.15 | mucin-associated surface protein (MASP, pseudogene) |
| Tc00.1047053508165.380 | -3.16 | hypothetical protein |
| Tc00.1047053509293.10 | -3.16 | mucin TcMUCII |
| Tc00.1047053508161.23 | -3.17 | hypothetical protein |
| Tc00.1047053509965.210 | -3.18 | hypothetical protein, conserved |
| Tc00.1047053511613.110 | -3.19 | mucin-associated surface protein (MASP) |
| Tc00.1047053510025.20 | -3.19 | mucin TcMUCII |
| Tc00.1047053510377.405 | -3.20 | hypothetical protein, conserved (pseudogene) |
| Tc00.1047053508389.138 | -3.21 | mucin TcMUCII |
| Tc00.1047053510023.10 | -3.22 | hypothetical protein, conserved (pseudogene) |
| Tc00.1047053506289.250 | -3.22 | surface protease GP63 (pseudogene) |
| Tc00.1047053506885.480 | -3.23 | trans-sialidase |
| Tc00.1047053511197.50 | -3.26 | elongation factor 1-gamma (EF-1-gamma, pseudogene) |
| Tc00.1047053508159.60 | -3.27 | mucin TcMUCII |
| Tc00.1047053506501.401 | -3.27 | hypothetical protein, conserved (pseudogene) |
| Tc00.1047053506973.25 | -3.29 | hypothetical protein |
| Tc00.1047053510025.210 | -3.33 | mucin TcMUCII |
| Tc00.1047053507957.43 | -3.36 | hypothetical protein, conserved (pseudogene) |
| Tc00.1047053511701.19 | -3.37 | surface protease GP63 |
| Tc00.1047053509847.70 | -3.38 | mucin TcMUCI |
| Tc00.1047053506667.40 | -3.39 | mucin-associated surface protein (MASP) |
| Tc00.1047053508839.70 | -3.44 | hypothetical protein, conserved |
| Tc00.1047053506359.80 | -3.48 | dynein light chain |
| Tc00.1047053511333.10 | -3.48 | calpain-like cysteine peptidase, Clan CA, family C2 |

| | | |
|---|---|---|
| Tc00.1047053506769.90 | -3.49 | surface protease GP63 |
| Tc00.1047053510377.61 | -3.50 | hypothetical protein, conserved |
| Tc00.1047053510275.410 | -3.50 | hypothetical protein |
| Tc00.1047053510279.50 | -3.50 | mucin TcMUCII (pseudogene) |
| Tc00.1047053511171.40 | -3.51 | trans-sialidase (pseudogene) |
| Tc00.1047053503577.9 | -3.54 | delta-1-pyrroline-5-carboxylate dehydrogenase (pseudogene) |
| Tc00.1047053511587.33 | -3.55 | hypothetical protein |
| Tc00.1047053508081.60 | -3.55 | hypothetical protein |
| Tc00.1047053506501.341 | -3.58 | surface protease GP63 (pseudogene) |
| Tc00.1047053511411.24 | -3.59 | hypothetical protein, conserved |
| Tc00.1047053505025.30 | -3.63 | mucin TcMUCII |
| Tc00.1047053508165.100 | -3.64 | mucin TcMUCII |
| Tc00.1047053506759.170 | -3.68 | mucin-associated surface protein (MASP) |
| Tc00.1047053511753.140 | -3.69 | vacuolar assembly protein vps41 |
| Tc00.1047053507981.20 | -3.70 | hypothetical protein |
| Tc00.1047053510841.30 | -3.70 | hypothetical protein |
| Tc00.1047053506599.70 | -3.72 | mucin TcMUCII |
| Tc00.1047053511797.167 | -3.78 | mucin-associated surface protein (MASP) |
| Tc00.1047053511603.420 | -3.84 | syntaxin binding protein (pseudogene), |
| Tc00.1047053507689.10 | -3.85 | dihydrouridine synthase (Dus, pseudogene) |
| Tc00.1047053408547.10 | -3.91 | mucin-associated surface protein (MASP) |
| Tc00.1047053511397.21 | -4.03 | elongation factor 1-gamma (EF-1-gamma, pseudogene) |
| Tc00.1047053510513.10 | -4.13 | mannosyl-oligosaccharide 1,2-alpha-mannosidase IB |
| Tc00.1047053506967.100 | -4.20 | mucin-associated surface protein (MASP) |
| Tc00.1047053510377.100 | -4.21 | mucin-associated surface protein (MASP) |
| Tc00.1047053506499.149 | -4.24 | mucin TcMUCII |
| Tc00.1047053503667.20 | -4.32 | hypothetical protein |
| Tc00.1047053511553.10 | -4.35 | mucin-associated surface protein (MASP) |
| Tc00.1047053509195.30 | -4.39 | mucin-associated surface protein (MASP) |
| Tc00.1047053509381.20 | -4.73 | 40S ribosomal protein S15A |
| Tc00.1047053506747.10 | -4.89 | (pseudogene),OMPDCase-OPRTase,point mutation |
| Tc00.1047053507237.190 | -5.02 | mucin-associated surface protein (MASP) |
| Tc00.1047053509201.20 | -5.08 | hypothetical protein, conserved |
| Tc00.1047053506731.10 | -5.51 | hypothetical protein |
| Tc00.1047053508541.225 | -8.74 | casein kinase |

Genes encoding cell surface proteins

**Supplementary Table 2. The number of differentially expressed genes and proteins across different sub-populations with a log2FC >2 and an adj. p-value <0.05.**

| Transcriptomics | Number of differentially expressed genes | Number of differentially expressed proteins | Proteomics |
|---|---|---|---|
| Epis - Amas | 195 | 583 | Epis - Amas |
| Epis - Trypos | 565 | 1309 | Epis - Trypos1 |
| | | 2124 | Epis - Trypos2 |
| Epis - QEpis | 291 | 470 | Epis - QEpi2 |
| | | 2342 | Epis - QEpis1 |
| Amas - Trypos | 596 | 1416 | Amas - Trypos1 |
| | | 1772 | Amas - Trypos2 |
| Amas - QEpis | 328 | 50 | Amas - QEpis2 |
| | | 1913 | Amas - QEpis1 |
| Trypos - QEpis | 833 | 1358 | Trypos1 - QEpis2 |
| | | 325 | Typos2 - QEpis1 |
| | | 1636 | Trypos1 - QEpis1 |
| | | 1761 | Trypos2 - QEpis2 |

**Supplementary Table 3. List of proteins present in QEpis 1 and 2, but absent from Epis (see also Fig. 4e)**

| Accession number | Group | Putative function |
|---|---|---|
| Q4D5A7_TRYCC | QEpi 1 and 2 | microtubule-associated protein |
| Q4CV65_TRYCC | QEpi 1 and 2 | RNA-binding protein 6 |
| Q4CPW8_TRYCC | QEpi 1 and 2 | trans-sialidase |
| Q4CR25_TRYCC | QEpi 1 and 2 | uncharacterized protein. |
| Q4DNY6_TRYCC | QEpi 1 and 2 | NUP-1 protein |
| Q4DNF9_TRYCC | QEpi 1 and 2 | trans-sialidase |
| Q4D335_TRYCC | QEpi 1 and 2 | dispersed gene family protein 1 (DGF-1) |
| Q4DLD0_TRYCC | QEpi 1 and 2 | trans-sialidase |
| Q4D749_TRYCC | QEpi 1 and 2 | ama-1 protein |
| Q4CNX7_TRYCC | QEpi 1 and 2 | SNARE protein |
| Q4DP71_TRYCC | QEpi 1 and 2 | uncharacterized protein. |
| Q4CTX4_TRYCC | QEpi 1 and 2 | trans-sialidase |
| Q4E649_TRYCC | QEpi 1 and 2 | uncharacterized protein. |
| Q4E0K2_TRYCC | QEpi 1 and 2 | proteasome regulatory ATPase subunit 3 |
| Q4D7C1_TRYCC | QEpi 1 and 2 | trans-sialidase. |
| Q4DGG5_TRYCC | QEpi 1 and 2 | vacuolar-type proton translocating pyrophosphatase 1 |
| Q4DA11_TRYCC | QEpi 1 and 2 | uncharacterized protein. |
| Q4D8Y8_TRYCC | QEpi 1 and 2 | nudix hydrolase domain-containing protein. |
| Q4CKC3_TRYCC | QEpi 1 | trans-sialidase |
| Q4DNH8_TRYCC | QEpi 1 | flagellum targeting protein kharon 1 |
| Q4D4F3_TRYCC | QEpi 2 | uncharacterized protein |
| Q4D532_TRYCC | QEpi 2 | QA-SNARE protein |
| Q4DFP4_TRYCC | QEpi 2 | uncharacterized protein |
